# Waking the sleepers: lincRNA overexpression compromises DHX36 activity and global protein synthesis

**DOI:** 10.64898/2026.06.28.735068

**Authors:** Robert Pasieka, Patrycja Plewka, Emanuele Vitale, Iga Kapuscinska, Mateusz Bajczyk, Dawid Bielewicz, Tomasz Skrzypczak, Kishor Gawade, Bartosz Koch, Agata Malinowska, Magdalena Bakun, Ewa Sitkiewicz, Alessia Ciarrocchi, Katarzyna Dorota Raczynska

**Affiliations:** Department of Gene Expression, Laboratory of RNA Processing, Institute of Molecular Biology and Biotechnology, Faculty of Biology, Adam Mickiewicz University, Poznan, Poland; Center for Advanced Technologies, Adam Mickiewicz University, Poznan 61-614, Poland; Laboratory of Translational Research, Azienda Unità Sanitaria Locale-IRCCS di Reggio Emilia, Reggio Emilia, Italy; Department of Gene Expression, Institute of Molecular Biology and Biotechnology, Adam Mickiewicz University, Poznan, Poland; Mass Spectrometry Laboratory, Institute of Biochemistry and Biophysics, Polish Academy of Sciences, 02-106 Warsaw, Poland

**Author notes:** Correspondence should be addressed to: Katarzyna Dorota Raczynska, Ph.D.

## Abstract

Transposable element-derived long intergenic noncoding RNAs (lincRNAs) are increasingly recognized as context-dependent regulators of gene expression; however, the functional consequences of their ectopic activation in somatic cells remain poorly understood. We previously showed that U7 snRNA represses a subset of LTR12-associated lincRNAs, including *lnc-ARRDC4-1* and *lnc-ADCYAP1-2*, two testis-enriched lincRNAs with minimal expression in somatic cells. Here, we examined the consequences of their increased expression in somatic cells. We showed that overexpression of either lincRNA led to common transcriptomic changes, proteomic changes, impaired migration, altered adhesion and proliferation, and a ∼50% reduction in protein synthesis. Furthermore, we identified *lnc-ARRDC4-1* as an upstream regulator of *lnc-ADCYAP1-2* transcription. Downstream of this event, *lnc-ADCYAP1-2* interacts with the RNA helicase DHX36, a regulator of G-quadruplex-containing mRNAs. *lnc-ADCYAP1-2* activation reduces DHX36 protein levels, which is accompanied by decreased protein output from a subset of DHX36 mRNA targets. At the cellular level, these effects correlate with cellular disfunction and altered global translation. Our results suggest a *lnc-ARRDC4-1:lnc-ADCYAP1-2*:DHX36 regulatory cascade linking the derepression of LTR12-containing lincRNAs to reduced protein synthesis and altered cellular processes in somatic cells.

## INTRODUCTION

Long intergenic noncoding RNAs (lincRNAs) constitute a major and functionally diverse class of the nonprotein-coding transcriptome. They are defined as RNA polymerase II transcripts exceeding 200 nucleotides in length whose genomic *loci* do not overlap with annotated protein-coding genes [1,2]. Although originally considered transcriptional noise, lincRNAs are now recognized as integral components of gene regulatory networks, with functions spanning both nuclear and cytoplasmic compartments [1]. In the nucleus, lincRNAs interact with chromatin-associated proteins, transcription factors, and regulatory complexes to modulate gene expression *in cis* at neighbouring *loci* or *in trans* at distant genomic regions. They also contribute to the formation and function of membraneless nuclear structures [3]. In the cytoplasm, lincRNAs regulate mRNA stability, modulate protein availability through decoy or scaffold functions, and influence translation efficiency [3–5]. Importantly, lincRNAs are increasingly viewed as interconnected components of regulatory networks rather than isolated transcriptional outputs. Within these networks, individual transcripts coordinate interactions across DNA, RNA, and protein layers to shape complex gene expression programs [3]. Moreover, a hallmark of lincRNAs is their pronounced cell type and tissue specificity, distinguishing them from most protein-coding transcripts and suggesting that they play roles in developmental and context-dependent regulatory programs rather than constitutive housekeeping functions [2].

More than two-thirds of human lincRNA genes contain sequences derived from transposable elements (TEs) [6,7]. Over evolutionary time, these elements have been coopted as functional promoters, enhancers, splice sites, and polyadenylation signals [6,7]. Among these, long terminal repeat 12 (LTR12) elements of human endogenous retroviral family 1 (HERV1) are enriched in CCAAT motifs recognized by the heterotrimeric transcription factor NF-Y [7,8]. Under physiological conditions, LTR12-driven transcription is tightly controlled and largely silenced in somatic tissues through multilayered epigenetic mechanisms, including DNA methylation and repressive histone modifications [9]. Consequently, LTR12 activity is predominantly restricted to germline tissues, particularly the testis, where it contributes to stage-specific transcriptional programs during spermatogenesis [10].

However, these latent somatic promoters can be reactivated under pathological or experimentally induced conditions. For example, inhibition of histone deacetylases induces widespread activation of LTR12-associated promoters across multiple somatic cell lines in an NF-Y-dependent manner, thereby altering gene expression programs linked to apoptosis and immune regulation [8]. Recent evidence further indicates that RNA-based mechanisms contribute to the somatic repression of these *loci*. In particular, U7 snRNA has been shown to repress specific LTR12 elements and LTR12-containing lincRNAs through direct base-pairing with HDE-like sequence motifs embedded within LTR12 sequences, which are short elements highly complementary to the 5′ end of U7 snRNA. This interaction attenuates the binding and/or transcriptional activity of NF-Y at adjacent CCAAT motifs, thereby maintaining transcriptional silencing of these *loci* in somatic cells [4]. In turn, depletion of U7 snRNA leads to the upregulation of numerous LTR12 elements together with a subset of LTR12-containing lincRNAs, among which *lnc-ARRDC4-1* and *lnc-ADCYAP1-2* are most prominently activated [4]. Both transcripts are robustly silenced in somatic cell lines, including HEK293T, HeLa, and SH-SY5Y cells and in somatic tissues; however, they can be preferentially expressed in testicular tissue, which is consistent with the germline-associated activity of LTR12 elements [4].

Here, we investigate the functional and molecular consequences of the ectopic activation of *lnc-ARRDC4-1* and *lnc-ADCYAP1-2* in human somatic cells. By integrating transcriptomic profiling, quantitative proteomics, and chromatin-associated RNA mapping, we identify a direct regulatory relationship between these two lincRNAs, in which *lnc-ARRDC4-1* promotes the transcriptional activation of *lnc-ADCYAP1-2* through its association with its genomic *locus*. Furthermore, we demonstrate that *lnc-ADCYAP1-2* interacts with DHX36 (DEAH box protein 36), a 3′−5′ RNA helicase with a strong affinity for resolving G-quadruplex (G4) structures in RNA [11–13]. The *lnc-ADCYAP1-2*:DHX36 interaction is accompanied by decreased protein output from a subset of DHX36 mRNA targets containing G4-forming sequences. At the cellular level, these effects correlate with altered cell proliferation, migration, adhesion, and global translation. Together, these findings reveal a regulatory axis linking the ectopic activation of germline-restricted lincRNAs to large-scale posttranscriptional rewiring through the modulation of DHX36.

## MATERIALS AND METHODS

### Cell culture and transfection

HEK293T cells (HEK WT, lnc-ARRDC4-1 OE and lnc-ADCYAP1-2 OE) were maintained in Dulbecco’s modified Eagle’s medium (Biowest) supplemented with 10% foetal bovine serum and antibiotics at 37°C in a humidified atmosphere containing 5% CO_2_, as previously described [4]. Stable cell lines overexpressing each long noncoding RNA (lncRNA) were generated as described previously [4]. Transient overexpression of *lnc-ARRDC4-1* and *lnc-ADCYAP1-2* was performed using plasmid-based transfection as described previously [4]. Cells were harvested 48–72 h posttransfection for downstream analyses.

### Cell proliferation assay

Cell proliferation was monitored in real time using an xCELLigence RTCA system (Agilent) with single E-plates for 72 h. The Cell Index values from 2-3 biological replicates were imported into R, interpolated to a common 15 min grid, and averaged. Doubling time was determined for the 10–72 h window as the first time point at which the Cell Index reached at least twice the starting value (CI at t = 10 h).

### Cell migration assay

Cell migration was assessed using the Oris™ Cell Migration Assay (Platypus Technologies, CMA1.101) according to the manufacturer’s protocol. Cells were seeded in 96-well plates with Oris™ Stoppers; after overnight attachment, the stoppers were removed to expose a 2 mm cell-free detection zone. Cells migrated for 48 h. Brightfield images were acquired at 0 h and 48 h using a Zeiss Axio Observer microscope, and detection zone coverage was quantified with ImageJ/Fiji. The detection zone was manually defined as a region of interest (ROI) in each image prior to quantification.

### Cell adhesion assay

Cells were seeded in µ-Dish 35 mm, high, glass-bottom dishes (Ibidi) and imaged live at 2 h postseeding. Thirty minutes prior to the 24 h imaging time point, the medium was replaced with medium containing Hoechst 33342 to label the cell nuclei for cell counting, and the cells were reimaged at 24 h. Cell adhesion and spreading were assessed by brightfield microscopy using a Zeiss Axio Observer. The cell area was manually measured at each time point using ImageJ/Fiji as a proxy for cell adhesion to the substrate.

### RNA isolation, reverse transcription and quantitative PCR

Total RNA was isolated using TRIzol reagent (Thermo Fisher Scientific) followed by TURBO DNase treatment or the Direct-zol RNA Miniprep Kit (Zymo Research), as previously described [4]. Reverse transcription was performed using random hexamers and SuperScript III (Thermo Fisher Scientific). Quantitative PCR was performed using SYBR Green chemistry (Applied Biosystems QuantStudio 6 Flex; 10 min at 95°C, then 40 cycles of 15 s at 95°C and 1 min at 60°C). Relative RNA levels were calculated using the ΔΔC_t_ method and normalized to those of *GAPDH* mRNA. The primer sequences are listed in Supplementary Table S1.

### Northern blot analysis

Northern blot analysis was performed as described previously [14], with modifications. 3 μg of total RNA was separated on a 1% denaturing agarose gel (1× HT buffer: 30 mM HEPES, 30 mM triethanolamine (pH 7.8), and 1.2% formaldehyde) at 100 V for 4 h with buffer recirculation. After mild alkaline treatment and neutralization, the RNA was transferred overnight to a Hybond N+ membrane in 10× SSC buffer and UV crosslinked (1200x100 µJ/cm^2^, UVP CL-1000S UV Crosslinker). Hybridization was performed with DIG-labelled DNA probes (DIG Oligonucleotide Tailing Kit, Roche), and detection was performed using an anti-digoxigenin antibody with CDP-Star substrate (Sigma‒Aldrich). Band intensities corresponding to individual pre-rRNA intermediates were quantified by densitometry using Image Studio (LI-COR Biosciences) software after background subtraction. Pre-rRNA processing was quantified using ratio analysis of multiple precursors (RAMPs). The ratios between selected pre-rRNA intermediates were calculated within each sample and used to assess relative changes in pre-rRNA processing.

### Subcellular fractionation

Cytoplasmic, nuclear, and chromatin-associated RNA fractions were isolated from lnc-ARRDC4-1 OE and lnc-ADCYAP1-2 OE cells using stepwise biochemical fractionation. All steps were performed on ice using LoBind tubes (Eppendorf) to maximize RNA recovery. Pelleted cells were resuspended in buffer A (10 mM HEPES (pH 7.5), 10 mM KCl, 1.5 mM MgCl₂, 0.1% NP-40, 0.5 mM DTT, 20% glycerol, 1× protease inhibitor cocktail (Roche), 40 U/ml ribonuclease inhibitor (Promega)), incubated for 5 min on ice, and centrifuged at 500 × g for 5 min at 4°C to yield the cytoplasmic fraction. The nuclear pellet was subsequently washed once with buffer B (20 mM HEPES (pH 7.5), 400 mM NaCl, 1.5 mM MgCl₂, 0.5 mM DTT, 20% glycerol, 1× protease inhibitor cocktail, 40 U/ml ribonuclease inhibitor), resuspended in buffer B, and incubated with buffer C (20 mM HEPES (pH 7.5), 150 mM NaCl, 3 mM MgCl₂, 1 mM DTT, 20% glycerol, 1× protease inhibitor cocktail, 40 U/ml ribonuclease inhibitor) for 30 min on ice. Centrifugation at 16000 × g for 10 min yielded the nuclear-soluble supernatant and the chromatin pellet. RNA was extracted from all fractions using TRIzol reagent and analysed by RT–qPCR. Fraction purity was validated by enrichment of *GAPDH* and *Actin* mRNAs in the cytoplasmic fraction and *MALAT1, KCNQ10T1,* and U1 snRNA in the nuclear/chromatin fractions, confirming minimal cross-contamination. Enrichment was expressed as log_2_ ratios between fractions, calculated from ΔΔCt values normalized to the corresponding input sample and averaged across three biological replicates (mean ± SD), with statistical significance assessed by two-sided Student’s *t*-test.

### Polysome profiling

Polysome profiling was performed in three biological replicates according to a protocol adapted from [15]. Total RNA was isolated from each of the four collected fractions (free subunits, 80S monosomes, light polysomes, and heavy polysomes) and reverse-transcribed, followed by qPCR for *DHX36* and *Actin* mRNAs. For each biological replicate, the relative amount of transcript in each fraction was calculated as 2^^(–Ct)^, and these values were normalized to the sum across all four fractions (Σ = 100%) to yield the percentage of total mRNA distributed to each fraction.

### Western blot analysis

Whole-cell protein extracts were analysed by SDS–PAGE and immunoblotting as previously described [4]. The membranes were probed with anti-DHX36 (Abcam, ab70269, 1:6000), anti-Actin (MP Biomedicals, 691001, 1:25000) or anti-β-Tubulin (Santa Cruz Biotechnology, sc-5274, 1:200) antibodies. Secondary antibodies conjugated with HRP included anti-mouse (Santa Cruz Biotechnology, sc-516102, 1:3000) and anti-rabbit (Santa Cruz Biotechnology, sc-2357, 1:3000) antibodies. Band intensities were quantified by densitometry using Image Studio and normalized to those of loading controls.

### SUnSET assay

Global protein synthesis was assessed using the SUnSET assay as previously described [16], with modifications. The cells were incubated with 5 μg/ml puromycin dihydrochloride (Gibco) for 15 min and then lysed in RIPA buffer. A total of 40 μg of protein was resolved on a 10% polyacrylamide gel and transferred to PVDF membranes (Millipore). The membranes were stained with Ponceau S and then probed with an anti-Puromycin antibody (Merck, clone 12D10, MABE343; 1:2000), followed by incubation with an anti-mouse secondary antibody (Santa Cruz, sc-516102; 1:2500). The signal was detected by enhanced chemiluminescence, quantified with Image Studio, and normalized to total protein (Ponceau S).

### RNA immunoprecipitation (RIP)

RIP was performed as described previously [4], with modifications to accommodate whole-cell lysates. Approximately 5×10⁶ HEK WT cells were washed twice with ice-cold PBS and lysed for 20 min on ice in RIP lysis buffer (20 mM Tris-HCl (pH 7.5), 150 mM NaCl, 1 mM MgCl₂, 0.5% NP-40, 1 mM DTT, 1× protease inhibitor cocktail, 40 U/ml ribonuclease inhibitor). The lysates were cleared by centrifugation at 12000 × g for 10 min at 4°C, and the supernatant was transferred to LoBind tubes. For each RIP reaction, 2 μg of anti-DHX36 antibody (Abcam, ab70269) or 2 μg of rabbit IgG (Santa Cruz Biotechnology, sc-2027, negative control) was incubated with 25 μl of Dynabeads Protein G (Invitrogen) for 1 h at 4°C with rotation. Antibody-bound beads were washed twice with lysis buffer and incubated with 500 μl of cleared lysate for 2 h at 4°C with gentle rotation. A total of 10% of each lysate was retained as input. Beads were washed four times with high-stringency RIP wash buffer (20 mM Tris-HCl (pH 7.5), 300 mM NaCl, 1 mM MgCl₂, 0.5% NP-40, 1 mM DTT, 40 U/ml ribonuclease inhibitor) and once with PBS. RNA was eluted by TRIzol extraction directly from the beads, followed by TURBO DNase treatment and ethanol precipitation. cDNA synthesis was performed using random hexamers and SuperScript III as described above. RIP signals were quantified by qPCR and are presented as a percentage of the input.

### Chromatin isolation by RNA purification and sequencing (ChIRP-seq)

ChIRP-seq was performed as described previously [17,18], with modifications. Briefly, 1×10⁸ lnc-ARRDC4-1 OE cells per biological replicate were crosslinked with 1% glutaraldehyde in PBS for 10 min at room temperature, quenched with 0.125 M glycine, and lysed in ChIRP lysis buffer (50 mM Tris-HCl (pH 7.0), 10 mM EDTA, and 1% SDS) supplemented with 100 U/ml Superase-In (Thermo Fisher Scientific) and 1× protease inhibitor cocktail. Chromatin was sheared by sonication using a Bioruptor Pico (Diagenode) to generate 200–500 bp fragments. A total of 1% of the sample was retained as input. Each sample was divided equally: half was hybridized with biotinylated antisense DNA probes targeting *lnc-ARRDC4-1,* whereas the remaining half was hybridized with biotinylated *LacZ* antisense probes used as a negative control (probe sequences are listed in Supplementary Table S2) [19]. The samples were diluted 1:3 in ChIRP hybridization buffer (750 mM NaCl, 1 mM EDTA, 1% SDS, and 15% formamide) and incubated overnight at 37°C with rotation. RNA–chromatin complexes were captured using Dynabeads MyOne Streptavidin C1 magnetic beads (Thermo Fisher Scientific) for 2 h at 37°C and then washed five times with ChIRP wash buffer (2× SSC, 0.5% SDS). DNA was purified from bead-bound complexes as described previously [18]. DNA libraries were prepared using the ThruPLEX® DNA-Seq Kit (Takara Bio, R400674) and sequenced on an Illumina NextSeq 500 platform using single-end 75 bp reads, as described previously [18].

Raw reads were quality checked with FastQC v0.11.8, trimmed with Trimmomatic v0.39, and aligned to the GRCh38/hg38 assembly using Bowtie2 v2.3.5.1. Duplicated and unmapped reads were removed using Picard v3.4.0 and SAMtools v1.9. Peak calling was performed with MACS2 v2.1.3.3 (*q*-value < 0.05, calculated using the Benjamini–Hochberg method) using *LacZ* pulldown as a background control. Peaks with MACS2 scores less than 125 were excluded from further analysis. Peak annotation was performed using the annotatePeak function from the ChIPseeker package with a transcription start site (TSS) window of ±3 kb relative to the GRCh38/hg38 annotations retrieved from Ensembl *via* AnnotationHub. Gene identifiers were verified against EnsDb, and gene names and biotypes were assigned using the org.Hs.eg.db database. All codes used for data processing and analysis are available on GitHub at https://github.com/Dawcio85/ChIRP_PLAB_Analysis.

### RNA sequencing and transcriptome analysis

RNA sequencing libraries were prepared from 1 μg of total RNA following rRNA depletion and sequenced as 100 bp single-end reads on an Illumina NextSeq 500/550 platform, as previously described [4]. Raw FASTQ files were quality-checked with FastQC v0.11.4 and adapter-trimmed with Cutadapt v3.7 (4). Trimmed reads were aligned to the GRCh38 human genome assembly using STAR v2.7.8a [20]. Raw count matrices were generated with featureCounts [21], and differential gene expression was analysed with DESeq2 [22]; genes with a Benjamini–Hochberg adjusted *p*-value (adj. *p*) < 0.05 were considered significant. Volcano plots were generated using ggplot2 [23].

### RNA antisense purification and mass spectrometry (RAP–MS)

RAP–MS was performed using UV-crosslinked HEK WT cells and biotinylated antisense DNA probes complementary to *lnc-ADCYAP1-2*. For UV crosslinking, the culture medium was removed, and the cells were washed twice with ice-cold PBS. Each 10 cm dish was overlaid with 2 ml of ice-cold PBS, placed on ice, and irradiated without the lid using 254 nm UV light at 150 mJ/cm² (UVP CL-1000S UV Crosslinker) to stabilize the RNA–protein interactions. The cells were scraped on ice, transferred to LoBind tubes, pelleted at 200 × g for 5 min at 4°C, washed once with cold PBS, and snap frozen in liquid nitrogen.

The cell pellets (∼1–1.5×10⁶ cells) were lysed in denaturing RAP lysis buffer (50 mM Tris HCl (pH 7.0), 10 mM EDTA, 1% SDS, 1 mM DTT, 1× protease inhibitor cocktail, and 80 U/ml ribonuclease inhibitor; 200 μl per sample) and incubated for 30 min on ice with occasional vortexing. The lysates were cleared by centrifugation at 16000 × g for 20 min at 4°C, and 5% of each lysate was saved as input for downstream RNA and MS analysis. For hybridization, cleared lysates were diluted 1:3 in RAP hybridization buffer (750 mM NaCl, 1 mM EDTA, 1% SDS, and 15% formamide) and incubated overnight at 37°C with biotinylated antisense probes targeting *lnc-ADCYAP1-2* (Lnc-ADCYAP2-1-ASO2 and Lnc-ADCYAP2-1-ASO5). Negative control reactions were hybridized with a control antisense probe (the probe sequences are listed in Supplementary Table S2). RNA–protein–probe complexes were captured using Dynabeads MyOne Streptavidin C1 (Invitrogen) preequilibrated in hybridization buffer. The beads were incubated with hybridized lysates for 2 h at 37°C with rotation and then washed five times with RAP wash buffer (2× SSC, 0.5% SDS) at 37°C to remove nonspecific interactions. For mass spectrometry sample preparation, on-bead digestion was performed by resuspending the beads in 50 μl of 50 mM ammonium bicarbonate containing 0.1% RapiGest and incubating them with sequencing grade trypsin (Promega) overnight at 37°C. The peptides were collected from the supernatant, acidified to hydrolyse RapiGest, cleared by centrifugation, and desalted using C18 StageTips.

The peptides were analysed by liquid chromatography–tandem mass spectrometry (LC–MS/MS) at the Mass Spectrometry Laboratory, Institute of Biochemistry and Biophysics, Polish Academy of Sciences (Warsaw, Poland). Depending on the analytical batch, measurements were performed using either an Orbitrap Exploris 480 or a Q Exactive mass spectrometer. The peptides were identified using the Mascot search engine (Matrix Science) against the UniProtKB/Swiss-Prot database (restricted to *Homo sapiens*). Searches were performed with trypsin specificity (up to 1 missed cleavage), methylthio (C) as a fixed modification and oxidation (M) as a variable modification. Monoisotopic mass values were used, with a precursor mass tolerance of ±5 ppm and a fragment mass tolerance of ±0.01 Da; protein mass was unrestricted. Peptide identification was accepted at a 1% false discovery rate (FDR). Protein abundance was estimated by the total number of MS/MS fragmentation spectra assigned to each protein. Statistical analysis of protein enrichment was performed using the IPinquiry4 R package (v0.4.7; https://github.com/hzuber67/IPinquiry4) [24]. Briefly, spectral counts were modelled using a Genewise Negative Binomial Generalized Linear Model implemented in edgeR, comparing *lnc-ADCYAP1-2*–specific probe pulldowns to control antisense probe pulldowns. Proteins with a log_2_-fold change (log_2_FC) > 0 and an adj. *p* < 0.05 were considered to indicate significant enrichment in the specific pull-down.

### DHX36 rescue experiment

HEK WT, lnc-ARRDC4-1 OE, and lnc-ADCYAP1-2 OE cells were seeded one day prior to transfection. Cells were transiently transfected using Lipofectamine 3000 (Thermo Fisher Scientific) with either an empty control pcDNA3.1(+) vector or a DHX36 overexpression vector. The DHX36 overexpression construct was generated by cloning the DHX36 coding sequence into the pcDNA3.1(+) backbone using the primers listed in Supplementary Table S1. Cells were harvested 72 h posttransfection and lysed as described above. Quantitative proteomic analysis was performed at the Mass Spectrometry Laboratory, Institute of Biochemistry and Biophysics, Polish Academy of Sciences (Warsaw, Poland) using a data-independent acquisition (DIA) workflow. Raw LC–MS/MS data were processed using DIA-NN against the human UniProt reference proteome and the cRAP contaminant database. Protein intensities were log_2_-transformed and normalized to the median signal intensity of each sample. Proteins detected in at least two samples in at least one experimental group were retained for downstream analyses, while contaminant proteins were removed. Missing values were imputed from a normal distribution. The statistical significance of differential protein abundance between experimental groups was assessed using Welch’s *t* test in Perseus, applying permutation-based FDR correction for multiple testing (*q*-value) < 0.05 or a *p-*value < 0.05, depending on the comparison. Principal component analysis (PCA) and volcano plots were generated for all pairwise comparisons. Gene Ontology enrichment analysis was performed using g:Profiler [25].

### R-loop data analysis

Published R-loop maps were obtained from R-loopBase (https://rloopbase.nju.edu.cn) [26], a database integrating 107 high-confidence genome-wide R-loop mapping datasets. Specifically, the Level 1 R-loop zone annotations were used, constituting a subset of the R-loop regions demonstrated by at least one experiment. ChIRP-seq peaks were intersected with Level 1 R-loop zones using the findOverlaps function from the GenomicRanges package v1.62.1 in R v4.5.2. A ChIRP-seq peak was classified as R-loop-associated if it overlapped with at least one Level 1 R-loop zone. The set of unique genes assigned to R-loop-overlapping ChIRP-seq peaks was subsequently intersected with the RNA-seq differential expression results to identify putative *lnc-ARRDC4-1* direct target genes regulated through R-loop-associated chromatin interactions. The cooccupancy of the ChIRP-seq signal with R-loop regions at selected *loci* was visualized using the Integrative Genomics Viewer (IGV).

### Triplex prediction

To identify candidate RNA:DNA triplex-forming regions within *lnc-ARRDC4-1* bound chromatin, genomic DNA sequences underlying ChIRP-seq peaks were extracted from the human reference genome (BSgenome.Hsapiens.UCSC.hg38, GRCh38/hg38) using the GenomicRanges and Biostrings packages in R. Because triplex-forming alignments can extend beyond the boundaries of a called peak, each peak was symmetrically extended by 300 bp on both flanks prior to sequence extraction; extensions were clipped where they would exceed chromosome boundaries. Triplex-forming potential between the repeat-masked *lnc-ARRDC4-1* transcript sequence (ENST00000562480.1) and each peak-derived DNA sequence was predicted using TriplexAligner [27], run in human mode. For each RNA:DNA:DNA sequence pair, only the best-scoring alignment (highest logE) was retained, and predicted triplexes were considered significant at E < 0.05 (logE > 1.301). Predicted triplex-forming alignments were required to overlap the coordinates of the originally called ChIRP-seq peak to be retained; alignments falling entirely within the flanking extension were excluded.

### Differential proteomic analysis

For all the quantitative proteomic experiments, the cells were washed once with warm PBS, harvested on ice using cold PBS and a cell scraper, and pelleted by centrifugation (200 × g, 5 min, 4°C). The cell pellets were snap-frozen in liquid nitrogen. Lysis buffer (50 mM Tris-HCl (pH 8.0), 150 mM NaCl, 1% SDS, and 1× protease inhibitor cocktail) was added to the frozen pellets, followed by 15 min of incubation on ice and 15 min of vortexing. The samples were boiled at 95°C for 10 min and sonicated for 15 min using a Bioruptor Plus [high-power, 30 s on/off cycles]; sonication was repeated if the lysate did not appear fully clarified. The lysates were cleared by centrifugation (16000 × g, 30 min, 4°C), and the supernatant was transferred to a new tube.

Quantitative proteomic analysis was performed at the Mass Spectrometry Laboratory, Institute of Biochemistry and Biophysics, Polish Academy of Sciences (Warsaw, Poland). For quantitative proteomic analysis, protein samples were subjected to tryptic digestion on filters using using Filter-Aided Sample Preparation (FASP) followed by tandem mass tag (TMT 10-plex) labeling and high pH fractionation into 12 fractions using Waters Acquity UPLC H-class system (Waters). Liquid chromatography–tandem mass spectrometry (LC–MS/MS) analysis was performed on a system comprising of Evosep One (Evosep Biosystems) coupled with Orbitrap Exploris 480 (Thermo Fisher Scientific) instrument. The raw data were processed using MaxQuant software (version 2.1). Protein identification was performed against the human UniProt reference proteome (version 2023_04) supplemented with popular contaminants. The statistical significance of differential protein abundance between experimental groups was assessed using Welch’s t-test in Perseus (version 1.6.15), applying permutation-based FDR correction for multiple testing, *q*-value < 0.05. PCA and volcano plots were generated for all pairwise comparisons. Gene Ontology enrichment analysis was performed using g:Profiler [25].

### AI use statement

Claude (Anthropic) was used to assist with code generation and refinement for data processing, statistical analysis and visualization. The authors directed, reviewed and validated all AI-assisted code and independently verified the resulting analyses.

### Data availability

The raw sequencing data generated in this study (RNA-seq and ChIRP-seq) will be publicly available upon publication. The code used for the ChIRP-seq data processing and analysis is available at https://github.com/Dawcio85/ChIRP_PLAB_Analysis. The mass spectrometry data will be available upon publication.

## RESULTS

### Overexpression of *lnc-ARRDC4-1* and *lnc-ADCYAP1-2* alters gene expression, protein synthesis and cell adhesion pathways

As shown in our previous report, silencing of U7 snRNA in HEK293T, HeLa, and SH-SY5Y cells leads to strong activation of two lincRNAs, *lnc-ARRDC4-1* and *lnc-ADCYAP1-2*, which are expressed at very low levels in the analysed somatic lines under wild-type conditions [4]. Interestingly, according to the GTEx data, their expression is limited primarily to testicular tissues (Supplementary Fig. S1) [4]. In this paper, we address the functional and molecular consequences of their increased levels in HEK293T cells.

To investigate the consequences of *lnc-ARRDC4-1* and *lnc-ADCYAP1-2* activation, we used previously generated HEK293T cell lines carrying CRISPR-mediated mutations within the HDE-like motifs of the respective *loci*, which disrupt U7-dependent repression and result in stable endogenous overexpression of the lincRNAs [4]. These cell lines, hereafter referred to as lnc-ARRDC4-1 OE and lnc-ADCYAP1-2 OE, were subjected to transcriptomic, proteomic and functional analyses.

RNA high-throughput sequencing (RNA-seq) revealed 2056 differentially expressed genes (DEGs) in lnc-ARRDC4-1 OE cells (1261 upregulated and 795 downregulated) and 1026 DEGs in lnc-ADCYAP1-2 OE cells (395 upregulated and 631 downregulated) (|log_2_FC| > 1 adj. *p* < 0.05) (Supplementary Tables S3 and S4; Supplementary Fig. S2A–C). A substantial fraction of these changes was shared between the two lines, with 234 common DEGs, 116 of which were upregulated and 118 of which were downregulated (Fig. 1A). GO biological process enrichment analysis revealed that among the upregulated transcripts, both lines were enriched for cell adhesion-related terms, cell motility and cell-cell signaling (Fig. 1B). The downregulated transcripts in both lines were enriched for neurogenesis and cell migration. Cell adhesion, embryo development and response to endogenous and abiotic stimuli were enriched specifically among the downregulated transcripts in lnc-ARRDC4-1 OE cells, whereas cell-cell adhesion, response to chemical and external stimuli, and tissue development were enriched among the downregulated transcripts in lnc-ADCYAP1-2 OE cells (Fig. 1C) [4].

**Figure 1.**
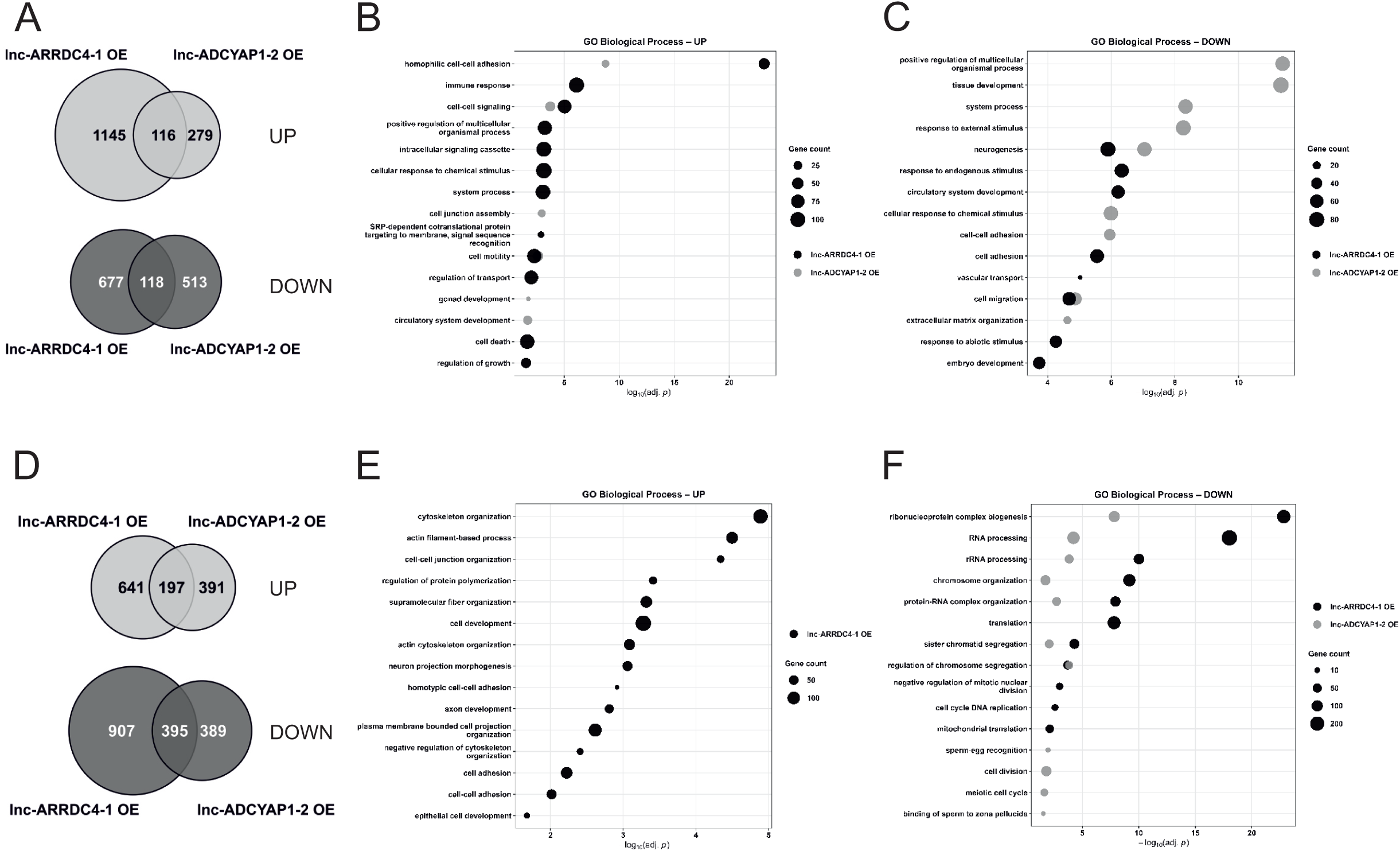
Overexpression of *lnc-ARRDC4-1* and *lnc-ADCYAP1-2* induces transcriptomic and proteomic changes. (A) Venn diagrams showing overlap between genes whose expression was upregulated (upper panel) and downregulated (lower panel) in lnc-ARRDC4-1 OE and lnc-ADCYAP1-2 OE cells. (B) GO biological process enrichment analysis of upregulated DEGs in lnc-ARRDC4-1 OE and lnc-ADCYAP1-2 OE cells. (C) GO biological process enrichment analysis of downregulated DEGs in lnc-ARRDC4-1 OE and lnc-ADCYAP1-2 OE cells. (D) Venn diagrams showing overlap between differentially expressed proteins (DEPs) upregulated (upper panel) and downregulated (lower panel) in lnc-ARRDC4-1 OE and lnc-ADCYAP1-2 OE cells. (E) GO biological process enrichment analysis of upregulated proteins in lnc-ARRDC4-1 OE cells. (F) GO biological process enrichment analysis of downregulated proteins in lnc-ARRDC4-1 OE and lnc-ADCYAP1-2 OE cells. For all the GO terms, fifteen representative nonredundant terms are shown. The dot size represents gene count; the x-axis shows −log_10_(adj. *p*). All the experiments were performed in three independent biological replicates. DEGs were defined as genes whose adj. *p* < 0.05 (calculated using the Benjamini–Hochberg method) and |log_2_FC| ≥ 1. DEPs were defined as proteins with a *q*-value < 0.01 (as calculated using Welch’s *t* test in Perseus and permutation-based FDR correction for multiple testing).

Furthermore, proteomic analysis revealed 2140 differentially expressed proteins (DEPs) in lnc-ARRDC4-1 OE cells (838 upregulated and 1302 downregulated) and 1372 DEPs in lnc-ADCYAP1-2 OE cells (588 upregulated and 784 downregulated) (*q*-value < 0.01) (Supplementary Tables S5 and S6; Supplementary Fig. S2D–F). A substantial fraction of these changes was shared between the two lines, with 592 common DEPs, of which 197 were upregulated and 395 were downregulated (Fig. 1D). Proteins whose level was increased in lnc-ARRDC4-1 OE cells were enriched for cytoskeleton organization, actin filament-based processes, cell–cell junction organization, and cell development, which is consistent with the observation that cytoskeletal remodelling accompanies altered cell adhesion and cell–cell adhesion (Fig. 1E). No significant GO enrichment was detected among the proteins whose level was increased in lnc-ADCYAP1-2 OE cells. Notably, proteins whose level was decreased in both lines were most significantly enriched for ribonucleoprotein complex biogenesis and organization, RNA and rRNA processing, and chromosome organization and segregation. Translation and DNA replication were also enriched among the downregulated proteins in lnc-ARRDC4-1 OE cells (Fig. 1F). In turn, the level of proteins related to cell division, the meiotic cell cycle, sperm-egg recognition and binding of sperm to zona pellucida were decreased in lnc-ADCYAP1-2 OE cells (Fig. 1E), which is consistent with the germline-associated expression pattern of this lincRNA (Supplementary Fig. S1).

Guided by both the overlapping transcriptomic and proteomic signatures in lnc-ARRDC4-1 OE and lnc-ADCYAP1-2 OE cells, we performed targeted functional assays. Consistent with the adhesion and motility GO terms, both lnc-ARRDC4-1 OE and lnc-ADCYAP1-2 OE cells exhibited impaired wound closure in the cell migration assay (Fig. 2A). Substrate adhesion was also decreased in lnc-ARRDC4-1 OE and lnc-ADCYAP1-2 OE cells, as evidenced by the reduced cell spreading area at 24 h postseeding compared with that of HEK WT cells (Fig. 2B). In addition, real-time impedance monitoring (xCELLigence) revealed reduced proliferation in both cell lines compared with that in HEK WT cells (Fig. 2C). Moreover, consistent with the proteomic changes related to translation and ribonucleoprotein complex biogenesis, compared with HEK WT cells, both lnc-ARRDC4-1 OE and lnc-ADCYAP1-2 OE cells showed an approximately 50% reduction in global protein synthesis, as assessed by the SUnSET assay, which measures puromycin incorporation into nascent polypeptides (Fig. 2D). Furthermore, to assess whether the proteomic enrichment of ribonucleoprotein complex biogenesis, including ribosome biogenesis and rRNA processing pathways reflects changes at the rRNA level, we examined pre-rRNA maturation by Northern blot using an ITS1 probe (Supplementary Fig. S3A). Both lnc-ARRDC4-1 OE and lnc-ADCYAP1-2 OE cells presented changes in pre-rRNA processing intermediate ratios compared with those of HEK WT cells, with reduced 41S/47S and 30S/47S ratios and increased 18S-E/47S, 18S-E/21S, and 18S-E/41S ratios (Fig. 2E; Supplementary Fig. S3), indicating that overexpression of either lincRNA affects pre-rRNA processing. In line with these findings, mature 18S, 5.8S, and 28S rRNA levels were significantly decreased, whereas 5S rRNA levels were increased in lnc-ARRDC4-1 OE cells. In the lnc-ADCYAP1-2 OE cells, 28S rRNA was also significantly reduced, while 18S, 5.8S, and 5S rRNA levels did not differ significantly from HEK WT (Fig. 2E).

**Figure 2.**
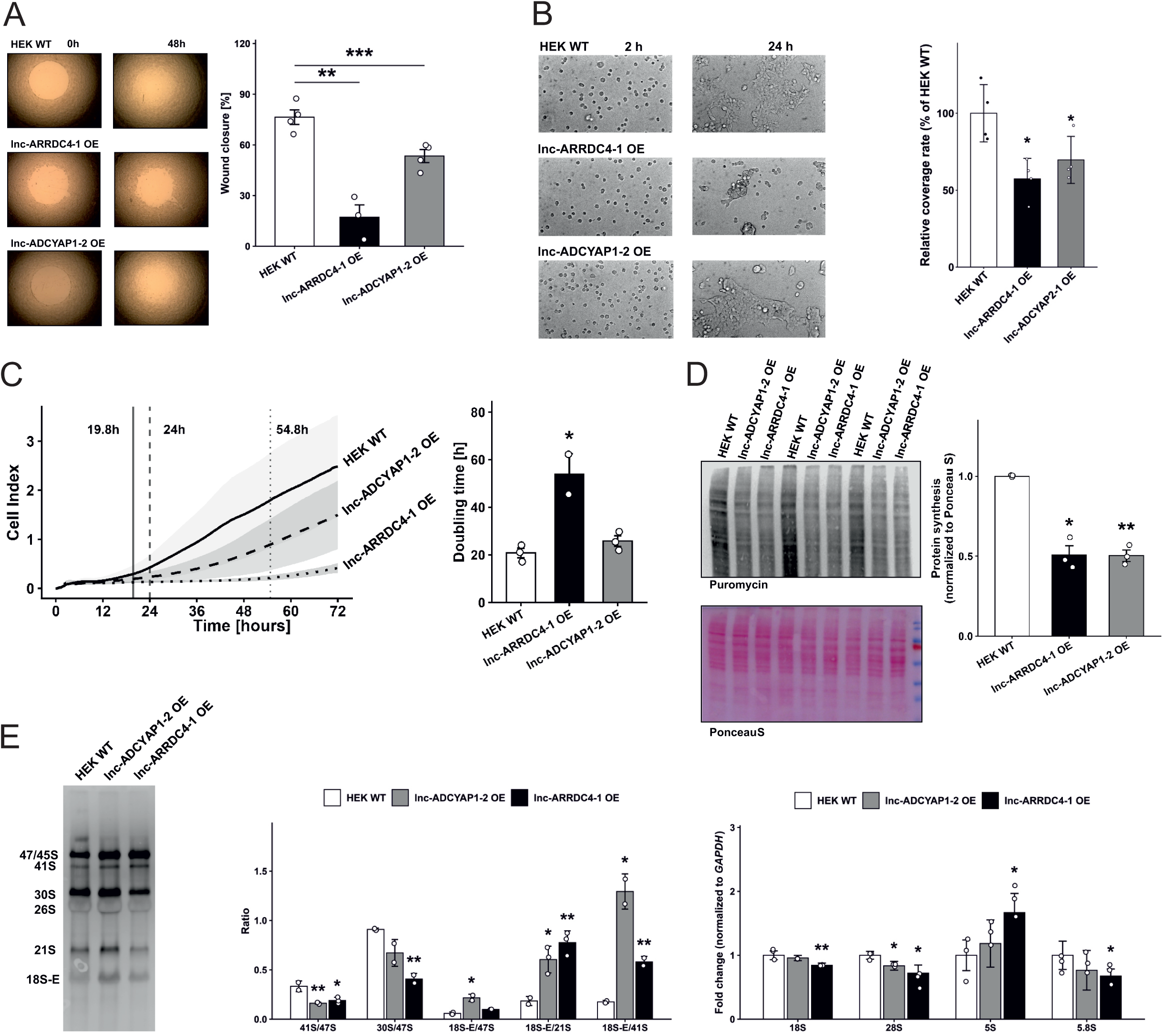
Overexpression of *lnc-ARRDC4-1* and *lnc-ADCYAP1-2* impairs cell proliferation, migration, adhesion, and protein synthesis. (A) Representative images of cell migration assays in HEK WT, lnc-ARRDC4-1 OE and lnc-ADCYAP1-2 OE cells at 0 h and 48 h after scratching (left panel). The wound closure quantification results are expressed as the percentage of the initial wound area (wight panel). (B) Representative phase-contrast images of HEK WT and lnc-ARRDC4-1 OE and lnc-ADCYAP1-2 OE cells at 2 h and 24 h after seeding. Scale bar, 100 μm (left panel). Quantification of the cell area at 24 h postseeding normalized to the cell area at 2 h postseeding in lnc-ARRDC4-1 OE and lnc-ADCYAP1-2 OE cells relative to HEK WT cells (right panel). (C) Real-time proliferation monitoring by an impedance-based xCELLigence assay in HEK WT, lnc-ARRDC4-1 OE and lnc-ADCYAP1-2 OE cells. Doubling times are indicated by vertical lines (left panel). Quantification of doubling times (right panel). (D) Representative Western blot of puromycin incorporation (SUnSET assay) and Ponceau S staining as a loading control in HEK WT, lnc-ADCYAP1-2 OE and lnc-ARRDC4-1 OE cells (left panel). Quantification of global protein synthesis normalized to the Ponceau S signal (right panel). (E) Northern blot analysis of pre-rRNA processing intermediates in HEK WT, lnc-ADCYAP1-2 OE and lnc-ARRDC4-1 OE cells using an ITS1 probe. The positions of the pre-rRNA intermediates (47/45S, 41S, 30S, 26S, 21S, and 18S-E) are indicated (left panel). Quantification of pre-rRNA intermediates ratios (41S/47S, 30S/47S, 18S-E/47S, 18S-E/21S, 18S-E/41S) in both overexpression lines relative to those in HEK WT cells (middle panel). Relative levels of mature rRNA species (18S, 28S, 5S, and 5.8S rRNA) normalized to *GAPDH* mRNA in HEK WT, lnc-ADCYAP1-2 OE and lnc-ARRDC4-1 OE cells, as determined by RT-qPCR (right panel). The data are presented as the mean ± SD from independent biological replicates (*n* = 4 for A; *n* = 3 for B, D, E; and *n* = 2–3 for C); the *p*-values were calculated using two-sided Student’s *t*-test (\**p*-value < 0.05; \*\**p*-value < 0.01; and \*\*\**p*-value < 0.001).

Overall, these transcriptomic, proteomic, and functional phenotypes across two independently generated cell lines strongly suggest that *lnc-ARRDC4-1* and *lnc-ADCYAP1-2* function through a shared regulatory pathway. Moreover, their activation in somatic cells has broad consequences for RNA metabolism, translational output, and cell behaviour.

### *lnc-ARRDC4-1* activates *lnc-ADCYAP1-2* transcription by associating with its *locus* through R-loop structure

To investigate the basis for the similar phenotypic effects observed in both lines, we first asked whether *lnc-ARRDC4-1* and *lnc-ADCYAP1-2* can regulate their own expression. We quantified *lnc-ARRDC4-1* levels in lnc-ADCYAP1-2 OE cells and, conversely, *lnc-ADCYAP1-2* levels in lnc-ARRDC4-1 OE cells by RT‒qPCR. These analyses revealed an approximately twofold increase in *lnc-ADCYAP1-2* transcript levels in lnc-ARRDC4-1 OE cells, with no significant effect on *lnc-ARRDC4-1* levels in lnc-ADCYAP1-2 OE cells, indicating that *lnc-ARRDC4-1* positively regulates *lnc-ADCYAP1-2* expression (Fig. 3A).

**Figure 3.**
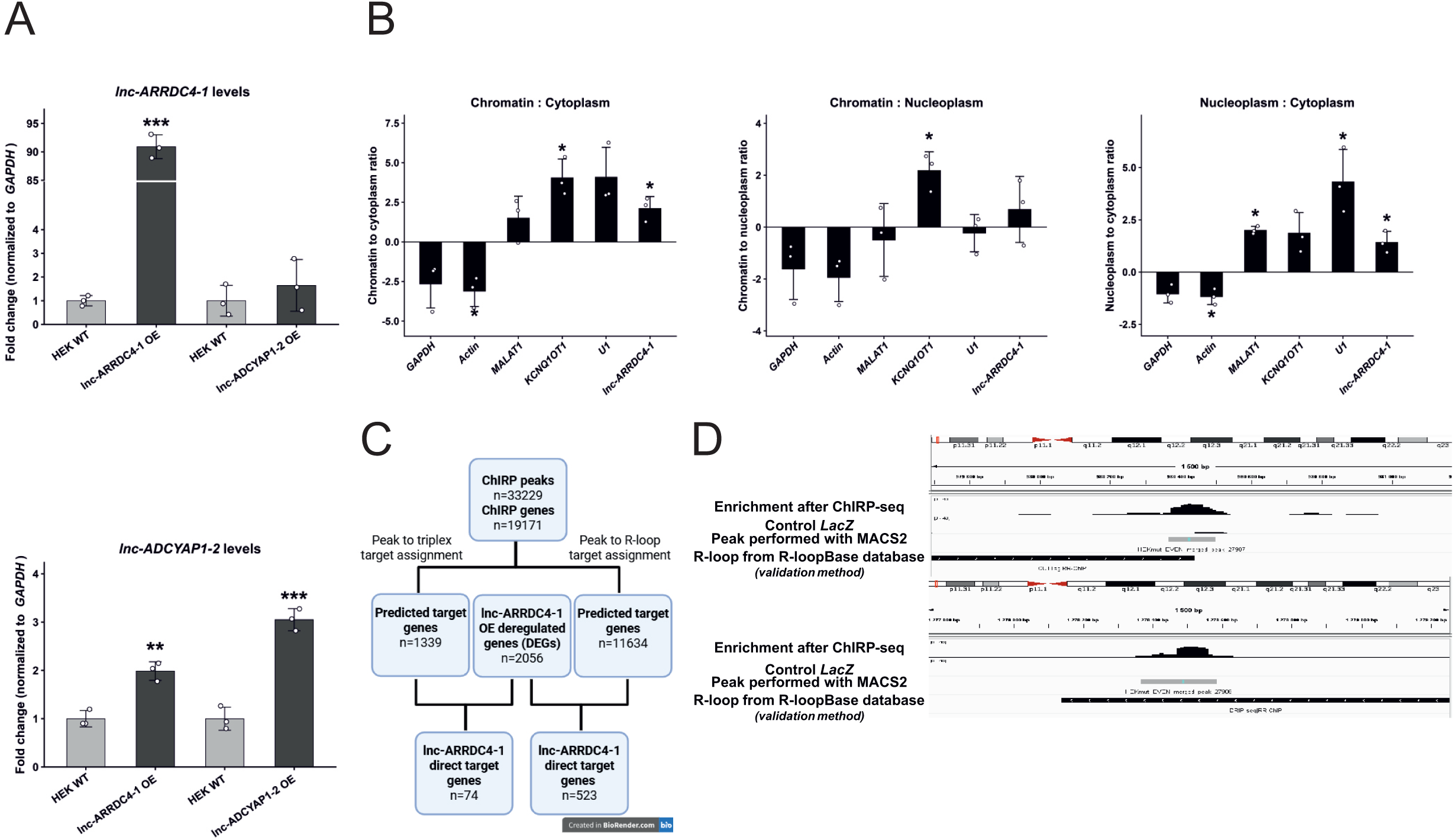
*lnc-ARRDC4-1* associates with the *lnc-ADCYAP1-2 locus* through R-loop formation and activates its transcription. (A) RT‒qPCR analysis of *lnc-ARRDC4-1* (upper panel) and *lnc-ADCYAP1-2* (lower panel) expression levels in lnc-ARRDC4-1 OE cells and lnc-ADCYAP1-2 OE cells relative to those in HEK WT cells. *GAPDH* levels were used for normalization. (B) Subcellular fractionation analysis of *lnc-ARRDC4-1* distribution across chromatin, nucleoplasmic, and cytoplasmic fractions in *lnc-ARRDC4-1* OE cells, shown as chromatin-to-cytoplasm, chromatin-to-nucleoplasm, and nucleoplasm-to-cytoplasm ratios. *GAPDH* and *Actin* were used as cytoplasmic controls; *MALAT1, KCNQ10T1,* and U1 snRNA were used as nuclear/chromatin controls. The data are presented as the mean ± SD (*n* = 3); the *p*-values were calculated using two-sided Student’s *t* test (\**p*-value < 0.05; \*\**p*-value < 0.01; and \*\*\**p*-value < 0.001). (C) Overlap of ChIRP-seq peaks with R-loop zones, predicted triplexes, and RNA-seq data. ChIRP-seq peaks (33229) assignment to target genes (19171) through two parallel approaches: overlap with predicted RNA–DNA triplex-forming regions using TriplexAligner (left branch) and overlap with experimentally mapped R-loop zones from R-loopBase (right branch), yielding 1339 and 11634 predicted target genes, respectively. Intersection of both target gene sets with lnc-ARRDC4-1 OE DEGs (2056) identified 74 triplex-based direct target genes and 523 R-loop-based direct target genes. (D) IGV genome browser views of two intronic regions of the *lnc-ADCYAP1-2 locus* at chr18:980,220–980,660 (upper) and chr18:1,278,062–1,278,942 (lower) covered with ChIRP-seq peaks, showing the *lnc-ARRDC4-1* ChIRP-seq signal, ChIRP-seq peak annotation, and Level 1 R-loop zones from R-loopBase. R-loop zones supported by experiments: CUTTag and RR-ChIP (upper) or DRIP-seq and RR-ChIP (lower).

Next, to determine the mechanism of *lnc-ADCYAP1-2* activation by *lnc-ARRDC4-1*, we first examined the subcellular localization of *lnc-ARRDC4-1* as a prerequisite for chromatin-based transcriptional control. Subcellular fractionation revealed that *lnc-ARRDC4-1* is predominantly nuclear and enriched in chromatin, with a positive chromatin-to-nucleoplasm ratio and a chromatin-to-cytoplasmic log_2_ ratio of 2.1, the latter comparable to those of the established chromatin-associated RNAs *MALAT1* and *KCNQ10T1* [28,29], and in stark contrast to cytoplasmic mRNAs, such as *GAPDH* and *Actin* (Fig. 3B). Such localization of *lnc-ARRDC4-1* suggests its potential role in transcriptional regulation at the *lnc-ADCYAP1-2 locus*.

Therefore, we decided to identify the global genomic *loci* occupied by *lnc-ARRDC4-1*. For this purpose, we performed ChIRP followed by high-throughput sequencing of DNA fragments (ChIRP-seq), which enables us to pull down chromatin fragments bound by *lnc-ARRDC4-1*. Enrichment of *lnc-ARRDC4-1* in ChIRP eluates relative to that in the *LacZ* controls was verified by RT–qPCR prior to library preparation and sequencing (Supplementary Fig. S4A). ChIRP-seq yielded 33229 high-confidence peaks belonging to 19171 genes (Supplementary Table S7; Supplementary Fig. S4B). Genomic annotation of these peaks revealed enrichment across three main categories: intronic regions (47.6% combined), distal intergenic regions (24.3%), and promoter-proximal regions within 3 kb of transcription start sites (21.1% combined) (Supplementary Table S7; Supplementary Fig. S4C). *lnc-ARRDC4-1* target gene distribution revealed enrichment of ChIRP peaks within lncRNAs (43.5%) and protein-coding genes (41.2%) (Supplementary Table S7; Supplementary Fig. S4D).

According to general knowledge, lncRNAs can associate with chromatin either by forming sequence-specific RNA:DNA:DNA triplex structures or by forming R-loops, in which the lncRNA hybridizes with complementary DNA strand and displaces the other strand as single-stranded DNA. To distinguish between these two modes of chromatin targeting, we cross-referenced ChIRP-seq peaks with RNA-seq data from lnc-ARRDC4-1 OE cells, computationally predicted triplex-forming sites [27], and experimentally mapped R-loop regions deposited in R-loopBase [26]. This analysis revealed 74 genes and 523 genes that may be directly regulated by *lnc-ARRDC4-1* through triplex-mediated chromatin association or through R-loop-mediated chromatin association, respectively (Fig. 3C), indicating that R-loop formation is the predominant mode through which *lnc-ARRDC4-1* associates with its chromatin targets.

Importantly, ChIRP-seq peaks also mapped to two intronic regions of the *lnc-ADCYAP1-2 locus*, colocalizing with independently mapped R-loop positions at both sites (Supplementary Table S7; Fig. 3D). Together, these data support a model in which *lnc-ARRDC4-1* occupies the *lnc-ADCYAP1-2* chromatin regions through R-loop formation, which in turn results in the activation of *lnc-ADCYAP1-2* transcription (Fig. 3A), providing a mechanistic explanation for the similar phenotypic effects observed in both lnc-ARRDC4-1 OE and lnc-ADCYAP1-2 OE cells (Fig. 2).

### *lnc-ADCYAP1-2* binds to DHX36 and impairs its translation

A substantial fraction of proteins altered in the lnc-ADCYAP1-2 OE cells were also changed in the lnc-ARRDC4-1 OE cells (∼50% of downregulated proteins (Fig. 1D, lower panel)), which is consistent with the finding that *lnc-ADCYAP1-2* acts as a downstream effector of *lnc-ARRDC4-1*. Both lines shared a proteomic phenotype characterized by enrichment of downregulated proteins involved in ribonucleoprotein complex biogenesis and organization, and rRNA processing (Fig. 1F). We therefore sought to identify the protein interaction partners of *lnc-ADCYAP1-2* that could account for this posttranscriptional perturbation and performed RNA antisense purification (RAP) followed by mass spectrometry. We identified DHX36 as one of the most prominent binding partners of *lnc-ADCYAP1-2*. Importantly, to strengthen our data, we conducted three different experimental RAP‒MS approaches: first, in cells with U7 snRNA knockdown, which increased endogenous *lnc-ADCYAP1-2* levels (Fig. 4A; Supplementary Table S8); second, in HEK WT cells (under physiological conditions) (Fig. 4B; Supplementary Table S9); and third, in HEK WT cells with transient overexpression of a plasmid encoding *lnc-ADCYAP1-2* (Fig. 4C; Supplementary Table S10). In all these analyses, DHX36 was identified as one of the most significantly enriched interactors. The *lnc-ADCYAP1-2*:DHX36 interaction was additionally confirmed by RIP using an anti-DHX36 antibody (Fig. 4D).

**Figure 4.**
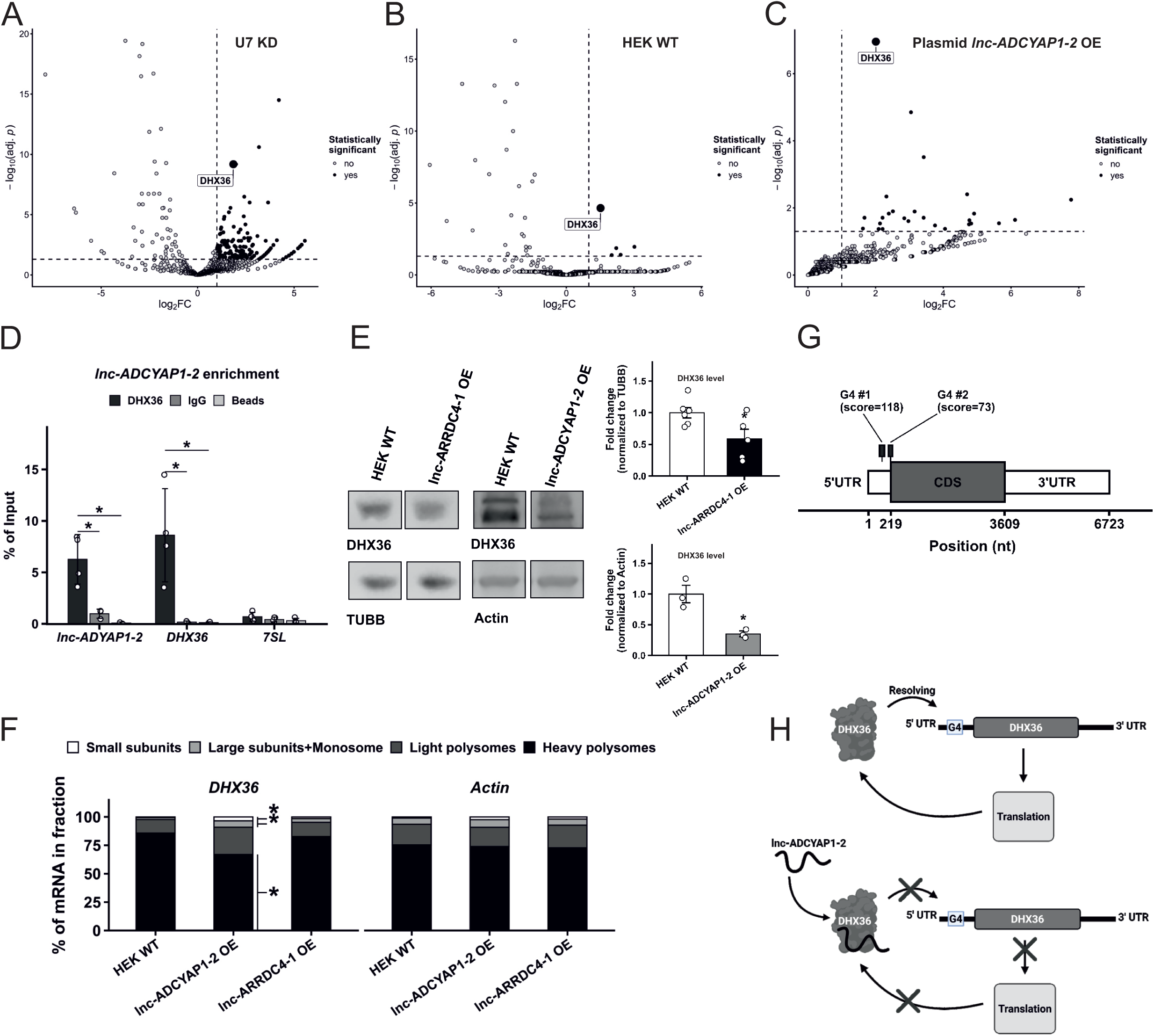
*lnc-ADCYAP1-2* binds DHX36 and reduces its levels. (A–C) Volcano plots of the RAP–MS data for *lnc-ADCYAP1-2*: (A) after U7 snRNA knockdown (U7 KD), which elevates endogenous *lnc-ADCYAP1-2* levels; (B) at the endogenous expression level in HEK WT cells; and (C) upon plasmid-driven overexpression. All the experiments were performed in three independent biological replicates. The dashed vertical and horizontal lines indicate the log_2_FC (log_2_FC > 1) and statistical significance (−log_10_(adj. *p*) > 1.3) thresholds, respectively. (D) RNA immunoprecipitation (RIP) followed by RT‒qPCR showing significant enrichment of *lnc-ADCYAP1-2* in the anti-DHX36-immunoprecipitated fraction relative to that in the IgG and bead control fractions in HEK WT cells. *DHX36* and the *7SL* served as positive and negative controls, respectively. (E) Representative Western blot of DHX36 and Actin protein levels in HEK WT, lnc-ARRDC4-1 OE and lnc-ADCYAP1-2 OE cells (left panel). Quantification of DHX36 protein levels normalized to that of Actin (right panel). (F) Polysome profiling of *DHX36* and *Actin* mRNAs in HEK WT, lnc-ADCYAP1-2 OE and lnc-ARRDC4-1 OE cells. The stacked bar charts show the distribution of mRNA across different fractions. (G) Schematic representation of the predicted G4 structures in *DHX36* mRNA. Two high-confidence G4 motifs (scores 118 and 73) are indicated within the 5′ UTR of the transcript. The positions are shown relative to the full-length *DHX36* mRNA (ENST00000496811.6). (H) Schematic representation of a proposed model in which *lnc-ADCYAP1-2* impairs DHX36 function. Under physiological conditions, the DHX36 protein resolves G4 structures within its own 5′ UTR, facilitating *DHX36* mRNA translation (upper panel). When *lnc-ADCYAP1-2* is overexpressed, *lnc-ADCYAP1-2* associates with DHX36, reducing its ability to unwind G4 structures. This, in turn, might impair the translation of *DHX36* mRNA, potentially resulting in decreased DHX36 protein levels. Data in D, E, and F are presented as the mean from independent biological replicates (*n* = 4 for D and *n* = 3 for E and F); *p*-values were calculated using two-sided Student’s *t*-test (\**p*-value < 0.05; \*\**p*-value < 0.01; and \*\*\**p*-value < 0.001).

Further analyses revealed that the protein level of DHX36 is reduced in lnc-ADCYAP1-2 OE and lnc-ARRDC4-1 OE cells. Quantitative proteomics revealed a modest decrease of DHX36 in both the lnc-ARRDC4-1 OE (|log2FC| = −0.08, *q*-value < 0.05) and lnc-ADCYAP1-2 OE datasets (|log_2_FC| = −0.48, *q*-value < 0.01; Supplementary Tables S3 and S4), which was further confirmed in Western blot analysis (relative level 0.59 and 0.36 in lnc-ARRDC4-1 OE and lnc-ADCYAP1-2 OE cells, respectively, vs. WT = 1; Fig. 4E). Importantly, the *DHX36* mRNA level remained unchanged under both conditions as determined by RT-qPCR and RNA-seq analyses (Supplementary Fig. S5; Supplementary Tables S3 and S4), indicating that the observed protein reduction occurred *via* a posttranscriptional mechanism.

To assess whether DHX36 translation is affected by lincRNA overexpression, we performed polysome profiling in HEK WT, lnc-ARRDC4-1 OE, and lnc-ADCYAP1-2 OE cells. We could not detect any changes in the global polysome profiles (Supplementary Fig. S6). However, as shown in Fig. 4F, when we isolated mRNAs from small subunits (40S), large subunits plus monosomes (60S and 80S), light polysomes and heavy polysomes and tested the distribution of *DHX36* transcripts by RT‒qPCR, we detected differences. In HEK WT cells, *DHX36* mRNA was predominantly associated with heavy polysomes, indicating active translation. In turn, in lnc-ADCYAP1-2 OE cells, the level of *DHX36* mRNA in the heavy polysome fraction was markedly reduced, with concomitant accumulation in lighter fractions, including light polysomes, large subunits, and monosome fractions, indicating impaired translational elongation and/or initiation. Notably, no significant redistribution was observed in the lnc-ARRDC4-1 OE cells (Fig. 4F). These results indicate that *lnc-ADCYAP1-2* specifically impairs *DHX36* translation, resulting in stronger reduction in the DHX36 protein level but no change in its mRNA level.

DHX36 is a DEAH-box RNA helicase with a strong affinity for G4 structures in RNA and has been shown to promote the translation of target mRNAs by unwinding inhibitory G4 structures within their 5′ UTRs [11,12]. To explore the basis for the observed translational inhibition of *DHX36* mRNA in lnc-ADCYAP1-2 OE cells, we examined whether the *DHX36* transcript itself contains G4 structures. *In silico* prediction using pqsfinder revealed two high-confidence G4 motifs within the *DHX36* transcript (scores 118 and 73), both mapping to the 5’ UTR (positions 108–157 and 195–236; Fig. 4G) [30]. Notably, *DHX36* mRNA itself was independently identified as a DHX36-bound target in a published PAR–CLIP dataset [11] and our RIP experiment (Fig. 4D), supporting the presence of functional G4 structures within this transcript and autoregulation pathway, which can potentially be disrupted by interaction with *lnc-ADCYAP1-2* (Fig. 4G and H).

### DHX36 targets are destabilized and less efficiently translated in lnc-ARRDC4-1 OE and lnc-ADCYAP1-2 OE cells

The reduction in the DHX36 protein level in lnc-ARRDC4-1 OE and lnc-ADCYAP1-2 OE cells raises the question of whether RNA targets of DHX36 are also affected at the proteomic level in these cells. To address this, we compared proteins whose levels were reduced in both overexpression lines against published PAR–CLIP data identifying mRNAs bound by DHX36 in human cells [11]. Interestingly, compared with 48.3% of all proteins detected in the MS dataset, 54.1% of the downregulated proteins in lnc-ARRDC4-1 OE cells and 54.5% of the downregulated proteins in lnc-ADCYAP1-2 OE cells corresponded to DHX36-bound mRNAs (*p*-value = 1.4×10⁻^6^ and *p*-value = 1.3×10⁻^4^, respectively; Fig. 5A). GO enrichment analysis of the DHX36 PAR–CLIP targets overlapping with reduced protein levels common to lnc-ARRDC4-1 OE and lnc-ADCYAP1-2 OE cells revealed ribonucleoprotein complex biogenesis and organization, RNA and rRNA processing, gene expression and biosynthetic processes as the most significantly enriched terms. Together with the effects on pre-rRNA processing (Fig. 2E), these findings might suggest the deregulation of ribosome assembly and function, further explaining the reduced translation observed in lnc-ARRDC4-1 OE and lnc-ADCYAP1-2 OE cells (Fig. 2D).

**Figure 5.**
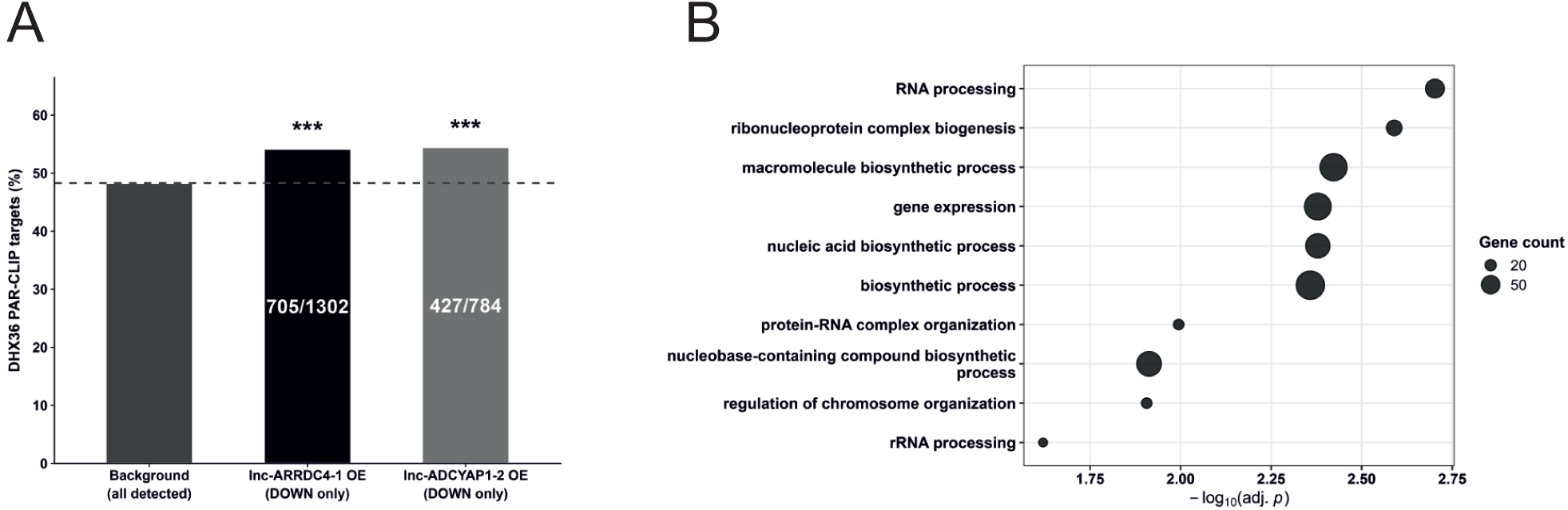
Translation of a set of proteins is affected by overexpression of *lnc-ARRDC4-1* and *lnc-ADCYAP1-2* and a reduced level of DHX36. (A) Enrichment of DHX36 PAR–CLIP targets among proteins whose level was reduced in lnc-ARRDC4-1 OE (54.1%) and lnc-ADCYAP1-2 OE cells (54.5%), shown as the percentage of overlapping DHX36 PAR–CLIP targets [11] compared with the background frequency (dashed line, 48.3%; all detected proteins; *p*-value < 0.05). (B) GO biological process enrichment analysis of all the downregulated DHX36 PAR–CLIP targets in both lnc-ARRDC4-1 OE and lnc-ADCYAP1-2 OE cells. The dot size represents gene count; the x-axis shows −log₁₀(adj. *p*).

In summary, we propose a model in which *lnc-ARRDC4-1* promotes the transcription of *lnc-ADCYAP1-2*, potentially through R-loop formation at its genomic *locus* (Supplementary Table S7; Fig. 3D). In turn, *lnc-ADCYAP1-2* binds to DHX36 (Supplementary Table S8–10; Fig. 4A–D), and this interaction reduces the DHX36 protein level posttranscriptionally, further impairing the translation of DHX36-bound mRNAs (Fig. 4E–H). Together, these data support a model in which two lincRNAs act in an axis to affect posttranscriptional gene regulation through DHX36.

To evaluate whether the proteomic changes induced by *lnc-ADCYAP1-2* overexpression are indeed linked to the reduction in DHX36 protein levels, we performed a rescue experiment in which we transiently overexpressed DHX36 in lnc-ADCYAP1-2 OE and lnc-ARRDC4-1 OE cells. To this end, HEK WT, lnc-ARRDC4-1 OE, and lnc-ADCYAP1-2 OE cells were first transiently transfected with an empty vector (EV, control), followed by quantitative proteomic analysis to identify proteins whose level was significantly altered by lncRNA overexpression. This analysis revealed 2276 DEPs in lnc-ARRDC4-1 OE cells and 1031 DEPs in lnc-ADCYAP1-2 OE cells (*q*-value < 0.05). Subsequently, lnc-ARRDC4-1 OE and lnc-ADCYAP1-2 OE cells were transfected with a DHX36 overexpression vector (DHX36 OE) (Supplementary Fig. S7A) and subjected to quantitative proteomic analysis. The resulting protein profiles were compared with those of the corresponding cells transfected with EV. Finally, these data were compared with the protein profiles of HEK WT cells transfected with EV to determine whether restoration of DHX36 expression in lnc-ARRDC4-1 OE and lnc-ADCYAP1-2 OE cells could reverse proteomic changes towards the wild-type cell state (Supplementary Tables S11 and S12).

Among the 2276 proteins with significantly altered levels in lnc-ARRDC4-1 OE cells, 211 (9.3%) were significantly increased in response to DHX36 overexpression (*p*-value < 0.05), of which 143 (67.8%) were restored to the wild-type level (Fig. 6A and 6B; Supplementary Tables S11; and Supplementary Fig. S7). In lnc-ADCYAP1-2 OE cells, 108 of 1031 proteins with altered levels (10.5%) responded to DHX36 overexpression, with 55 (50.9%) restored to the wild-type level (Fig. 6A and 6B; Supplementary Tables S12; and Supplementary Fig. S7). These results indicate that DHX36 overexpression in lnc-ARRDC4-1 OE and lnc-ADCYAP1-2 OE cells partially reverses the proteomic changes induced by decreased DHX36 level due to lincRNA overexpression, supporting a functional link between reduced DHX36 activity and the observed proteomic phenotype.

**Figure 6.**
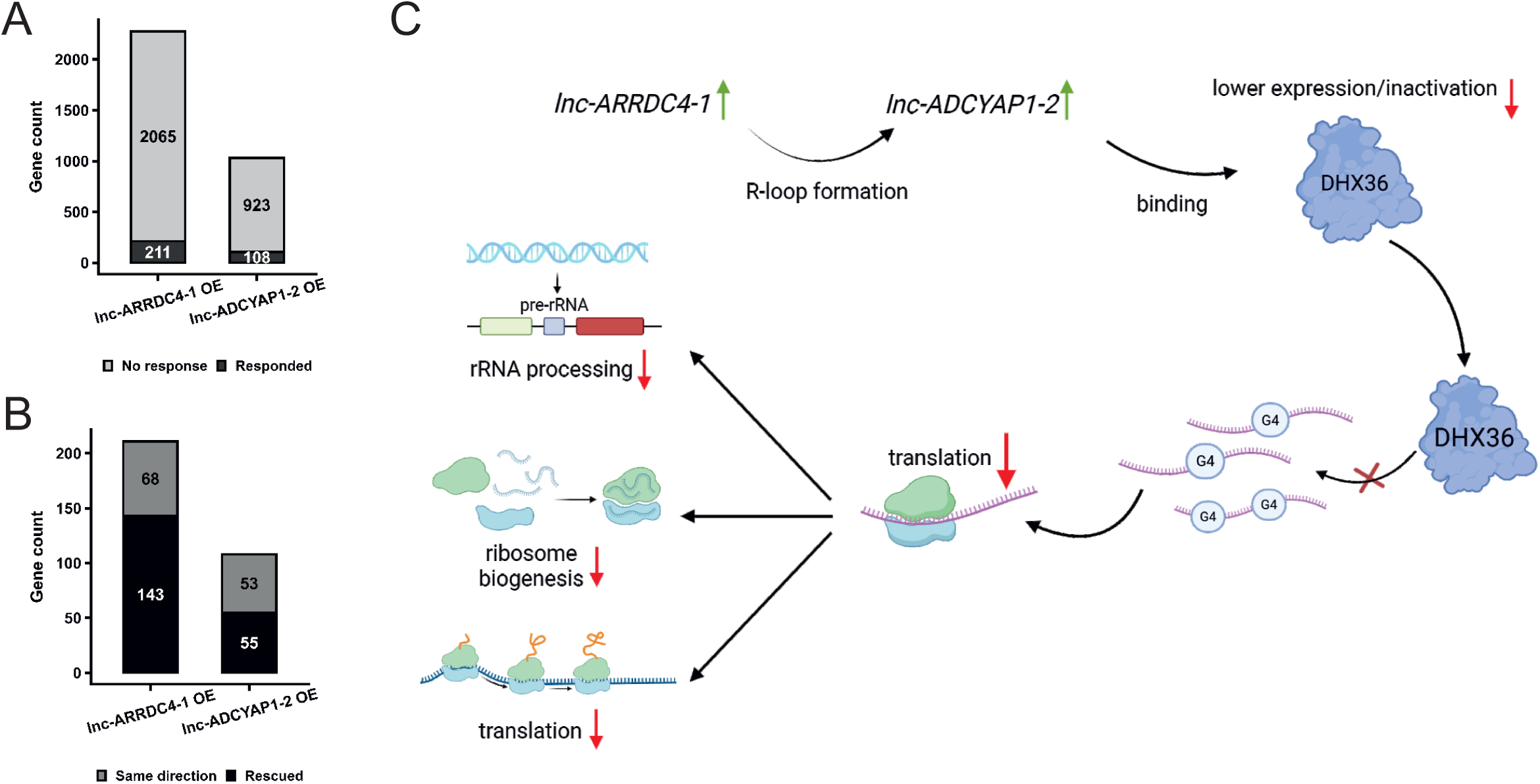
DHX36 overexpression partially rescues the proteomic changes induced by *lnc-ARRDC4-1* and *lnc-ADCYAP1-2* activity. (A) Stacked bar charts showing the total number of proteins with altered levels in lnc-ARRDC4-1 OE and lnc-ADCYAP1-2 OE cells relative to that in HEK WT cells transfected with the empty vector, stratified by response to DHX36 overexpression: the dark grey bars indicate the proteins whose level significantly changed *p*-value < 0.05) upon overexpression of DHX36; the light grey bars indicate the proteins whose level did not significantly respond. This experiment was performed in three independent biological replicates. (B) Stacked bar charts showing the direction of changes among proteins that responded to DHX36 overexpression (shown in A): the black bars represent the proteins restored to levels observed in HEK WT cells transfected with EV; the grey bars represent proteins whose level changed in the same direction as that under lncRNA overexpression conditions. (C) Proposed model of the *lnc-ARRDC4-1*:*lnc-ADCYAP1-2*:DHX36 regulatory axis. *lnc-ARRDC4-1* upregulation drives the transcriptional activation of *lnc-ADCYAP1-2 via* R-loop formation at the *lnc-ADCYAP1-2 locus*, resulting in its accumulation. In turn, *lnc-ADCYAP1-2* binds to the DHX36 protein, leading to reduced DHX36 levels and function. DHX36 loss impairs the translation of its target mRNAs, ultimately resulting in decreased rRNA processing, disrupted ribosome biogenesis and translation.

## DISCUSSION

### *lnc-ARRDC4-1* and *lnc-ADCYAP1-2* form a hierarchical regulatory axis downstream of U7 snRNA-mediated repression

In this report, we show that two LTR12-containing lincRNAs, *lnc-ARRDC4-1* and *lnc-ADCYAP1-2*, which are maintained in a repressed state in somatic cells partially by U7 snRNA [4], function within a hierarchical regulatory cascade when U7 snRNA-mediated repression is relieved. *lnc-ARRDC4-1* occupies the *lnc-ADCYAP1-2* genomic *locus* and promotes its transcription, potentially through R-loop formation at its genomic region. The *lnc-ADCYAP1-2* transcript further binds to the RNA helicase DHX36 and interferes with DHX36-dependent translation, reducing the protein output of a subset of DHX36 targets. This cascade is associated with the deregulation of the levels of a significant number of proteins and disturbances in RNA processing, rRNA processing and ribonucleoprotein complex biogenesis, including ribosome biogenesis (Fig. 6C). These results reveal a downstream consequence of U7-mediated lincRNA derepression and connect LTR12-derived lincRNAs with DHX36-dependent posttranscriptional regulation.

The ability of *lnc-ARRDC4-1* to activate *lnc-ADCYAP1-2* transcription is consistent with previous reports describing hierarchical lncRNA regulatory cascades, in which one lncRNA directly controls the expression of another noncoding transcript through chromatin-associated mechanisms [3,31]. Indeed, *lnc-ARRDC4-1* ChIRP-seq analysis identified 33229 high-confidence peaks, with a binding pattern enriched at promoter-proximal and intronic regions (Supplementary Fig. S4C). Furthermore, the colocalization of *lnc-ARRDC4-1* ChIRP-seq peaks with independently mapped R-loop zones at two intronic sites within the *lnc-ADCYAP1-2 locus* (Fig. 3D) suggests that *lnc-ARRDC4-1* may act *in trans* at this region, forming an R-loop through direct hybridization with the complementary DNA strand. A comparable *trans* R-loop-associated targeting mechanism has recently been reported for the lncRNA *TERRA*, which was shown to bind nontelomeric genomic *loci in trans* and form R-loops at these sites through direct RNA:DNA hybridization [32]. The overlap between the *lnc-ARRDC4-1* ChIRP-seq peaks and publicly available R-loop mapping data at the *lnc-ADCYAP1-2 locus* is consistent with this interpretation. Nevertheless, further experimental evidence is needed to confirm this mechanism, including experiments designed to disrupt the RNA:DNA hybrid between the *lnc-ARRDC4-1* transcript and the *lnc-ADCYAP1-2 locus*, for example, through RNase H overexpression or targeted mutation of the hybridization-competent sequence.

### *lnc-ADCYAP1-2* reduces DHX36 protein levels and activity which in turn affects global protein synthesis

DHX36 is a predominantly cytoplasmic DEAH-box RNA helicase with a well-characterized preference for resolving G4 structures in RNA [11,13]. PAR–CLIP mapping revealed that DHX36 binds thousands of mRNA targets enriched in G4-forming sequences within their 5′ UTRs and promotes their translational efficiency; loss of DHX36 leads to the accumulation of these transcripts in translationally inactive states [11]. *lnc-ADCYAP1-2* is partially localized in the cytoplasm (Supplementary Fig. S8), where it can physically associate with DHX36, as shown by RAP–MS under three independent conditions: i) following U7 snRNA knockdown (Fig. 4A; Supplementary Table S8); ii) at endogenous *lnc-ADCYAP1-2* expression levels in HEK WT cells (Fig. 4B; Supplementary Table S9); and iii) following transient overexpression of *lnc-ADCYAP1-2* from a plasmid (Fig. 4C; Supplementary Table S10), and confirmed by RIP (Fig. 4D). Whether this association reflects direct binding or occurs within a larger ribonucleoprotein complex remains to be determined. Whether this association reflects direct binding or occurs within a larger ribonucleoprotein complex remains to be determined. A conceptually similar regulatory scheme has been described for the lncRNA *GSEC*. *GSEC* binds DHX36 and antagonizes its G-quadruplex-resolving activity, thereby promoting colon cancer cell migration [33].

The overexpression of *lnc-ARRDC4-1* and *lnc-ADCYAP1-2* reduces the DHX36 protein level without affecting the *DHX36* transcript level, and polysome profiling revealed a marked shift in the *DHX36* mRNA level from the heavy polysome fraction to the light polysome fraction in lnc-ADCYAP1-2 OE cells (Fig. 4E and F), indicating that translational attenuation was the primary mechanism. The absence of this effect in lnc-ARRDC4-1 OE cells may be attributed to the less pronounced reduction in DHX36 protein levels observed in these cells (Fig. 4E, Supplementary Table S5). This finding is also consistent with an indirect mechanism mediated by the induction of *lnc-ADCYAP1-2*, rather than direct regulation of the helicase.

The *DHX36* transcript harbours two predicted high-confidence G4 motifs within its 5′ UTR (scores of 118 and 73 according to pqsfinder; Fig. 4G), and the DHX36 protein has been shown to bind its own mRNA in PAR–CLIP [11] and RIP (Fig. 4D) experiments. DHX36 promotes the translation of target mRNAs by unwinding 5′ UTR G4 structures [11,13]. These observations suggest that DHX36 may also facilitate its own translation through G4 unwinding within the *DHX36* 5′ UTR. One possibility is that the binding of *lnc-ADCYAP1-2* interferes with this process, thereby reducing the DHX36 protein output (Fig. 4H). This model requires direct experimental validation, for example, using luciferase reporter constructs containing wild-type and G4 motif-mutated *DHX36* 5′ UTRs expressed under conditions with or without DHX36 expression.

The reduction in global protein synthesis in both lnc-ARRDC4-1 OE and lnc-ADCYAP1-2 OE cells—approximately 50% relative to that in HEK WT cells, as measured by the SUnSET assay (Fig. 2D)—is consistent with the proteomic changes observed in these cells and the reduced translation of DHX36 targets. Proteins whose level was reduced in both cell types were most strongly enriched for ribonucleoprotein complex biogenesis and organization, rRNA processing and translation (Fig. 1F), processes that underpin ribosome function and translational capacity. Proteins involved in the same processes were also enriched among DHX36 PAR–CLIP targets, which were overrepresented among the downregulated proteins in lnc-ARRDC4-1 OE and lnc-ADCYAP1-2 OE cells (Fig. 5A). The partial rescue of proteomic changes by transient overexpression of DHX36 (Fig. 6A and B) supports the direct contribution of reduced DHX36 activity to this phenotype, although the incomplete nature of the rescue indicates that additional mechanisms might contribute.

Furthermore, the observed translational impairment is consistent with the reduced levels of 18S, 5.8S, and 28S rRNAs in lnc-ARRDC4-1 OE cells and to a lesser extent in lnc-ADCYAP1-2 OE cells, providing further evidence for a broader defect in ribosome biogenesis affecting both ribosomal subunits (Fig. 2E). Notably, the increase in 5S rRNA in the lnc-ARRDC4-1 OE cells, together with decreased levels of the other rRNAs constituting the 60S subunit, may point to an imbalance between 5S rRNA production and large subunit maturation. The altered levels of mature rRNAs may be partially explained by defects in rRNA maturation. For instance, the accumulation of the 18S-E precursor may reflect impaired processing into mature 18S rRNA. Consistent with this, our quantitative proteomic data revealed reduced levels of NOB1, RIOK1, and EIF1AD, proteins involved in the final stages of pre-40S subunit maturation and 18S-E processing [34,35] (Supplementary Table S5 and S6). Although the observed alterations in rRNA levels do not appear to substantially affect polysome assembly, as evidenced by the absence of significant changes in polysome profiles (Supplementary Fig. S6), they may nevertheless impair global translational efficiency by affecting ribosome activity and the rates of translation initiation and elongation. This possibility is further supported by the reduced levels of translation initiation and elongation factors observed in both lnc-ARRDC4-1 OE and lnc-ADCYAP1-2 OE cells (Supplementary Tables S5 and S6).

### Biological relevance: A germline-specific regulatory axis ectopically activated in somatic cells

Both *lnc-ARRDC4-1* and *lnc-ADCYAP1-2* are highly restricted in normal human tissues, with detectable levels predominantly in the testis according to GTEx data (Supplementary Fig. S1) [4]. This pattern reflects the known biology of their LTR12 elements, which are subjected to NF-Y-driven transcriptional activation specifically in spermatogonial stem cells and early differentiated spermatogonia and are otherwise maintained in a silenced state in somatic tissues through DNA methylation, repressive histone modifications, and U7 snRNA-mediated restriction [4,9,36]. The regulatory axis described here may therefore represent a component of the gene expression program operating during early spermatogenesis, where widespread transcription from transposable element-derived promoters has been documented [37,38]. However, when reactivated in somatic cells, as shown here, they may resemble, at least in part, a germline-specific gene expression program in a somatic context, with potential functional consequences, including impaired RNA metabolism and translation along with altered cell adhesion and migration.

### Limitations

Several aspects of the proposed model require further experimental support. The colocalization of *lnc-ARRDC4-1* ChIRP-seq peaks with R-loop zones at the lnc-*ADCYAP1-2 locus* supports transcriptional activation; however, whether R-loop formation is indeed required for *lnc-ADCYAP1-2* activation remains to be experimentally tested. The molecular basis of the *lnc-ADCYAP1-2*:DHX36 interaction—whether it involves specific structural motifs within *lnc-ADCYAP1-2*, direct competition with G4-containing mRNA substrates, or another mechanism—remains to be determined. The results of rescue experiments revealed that DHX36 loss accounts for only a fraction of the proteomic changes induced by lincRNA overexpression, suggesting the involvement of additional effector mechanisms. Finally, all the experiments were performed in HEK293T cells; whether the regulatory axis operates similarly in other cell types or in the physiologically relevant context of testicular germ cells requires further investigation.

## Acknowledgements

We thank Prof. Eliza Wyszko and Agata Tyczewska, Ph.D., for providing access to the xCELLigence system and Marta Orlicka-Płocka, Ph.D., and Dorota Gurda-Woźna, Ph.D., for their technical assistance with the real-time proliferation monitoring by the impedance-based xCELLigence system. We thank Markus Hafner, Ph.D., for discussing his published data.

## Author contributions

R.P. conceived the study, designed and performed the experiments, analysed the data, prepared all the figures and tables, and wrote the manuscript. P.P. generated stable CRISPR-Cas9-mutant cell lines; performed the RT‒qPCR (Fig. 3A), growth curve, and RAP‒MS experiments; and contributed to the experimental planning, data interpretation, and manuscript editing. E.V. assisted in performing the ChIRP experiment, prepared the DNA sequencing libraries, and performed the DNA sequencing. I.K. performed the SUnSET experiment, polysome profiling, and Northern blot analysis and edited the manuscript. M.B. performed the bioinformatic processing of the RAP–MS data and edited the manuscript. D.B. performed the bioinformatic processing of ChIRP-seq data and edited the manuscript. T.S. performed the fluorescence microscopy imaging and edited the manuscript. K.G. performed the bioinformatic processing of the RNA-seq data and edited the manuscript. B.K. prepared the samples for differential proteomics and edited the manuscript. A.M. performed proteomics experiment and analysis. M.B. performed proteomics experiment and analysis. E.S. performed proteomics experiment and analysis. A.C. contributed to the concept and planning of the ChIRP-seq experiment and edited the manuscript. K.D.R. conceived the study, designed the experiments, interpreted the results, and edited the figures and the manuscript.

---

**Supplementary Figure S1. Tissue expression profiles of *lnc-ARRDC4-1* and *lnc-ADCYAP1-2*.** Bulk tissue gene expression profiles (TPM) of *lnc-ARRDC4-1* (ENSG00000259870; upper panel) and *lnc-ADCYAP1-2* (ENSG00000263551; lower panel) from the GTEx database. Each dot represents a single sample; the box plots indicate the median and interquartile range per tissue.

**Supplementary Figure S2. Transcriptomic and proteomic overview of lnc-ARRDC4-1 OE and lnc-ADCYAP1-2 OE cells.** (A) PCA of RNA-seq data from HEK WT, lnc-ARRDC4-1 OE and lnc-ADCYAP1-2 OE cells. (B) Volcano plot of DEGs in lnc-ARRDC4-1 OE cells relative to those in HEK WT cells. (C) Volcano plot of DEGs in lnc-ADCYAP1-2 OE cells relative to those in HEK WT cells. The numbers indicate the number of genes that were significantly upregulated (right) or downregulated (left). (D) PCA of MS data from HEK WT, lnc-ARRDC4-1 OE and lnc-ADCYAP1-2 OE cells. (E) Volcano plot of DEPs in lnc-ARRDC4-1 OE cells relative to those in HEK WT cells. (F) Volcano plot of DEPs in lnc-ADCYAP1-2 OE cells relative to those in HEK WT cells. The numbers indicate the number of proteins that were significantly upregulated (right) or downregulated (left). All the experiments were performed in three independent biological replicates. DEGs were defined as genes with an adj. *p* < 0.05 (calculated using the Benjamini–Hochberg method) and |log_2_FC| ≥ 1. DEPs were defined as proteins with a *q*-value < 0.01 (as calculated using Welch’s *t*-test in Perseus and permutation-based FDR correction for multiple testing).

**Supplementary Figure S3. Northern blot analysis of pre-rRNA processing intermediates.** (A) Scheme showing the 47S pre-rRNA with the positions of the probe used in the Northern blot and rRNA biogenesis pathway analyses. (B) Ethidium bromide (EtBr)-stained agarose gel showing total RNA from HEK WT, lnc-ADCYAP1-2 OE and lnc-ARRDC4-1 OE cells, used as a loading control. (C) Northern blot hybridization with an ITS1 probe to detect pre-rRNA intermediates (47/45S, 41S, 30S, 26S, 21S and 18S-E) in HEK WT, lnc-ADCYAP1-2 OE and lnc-ARRDC4-1 OE cells. This experiment was performed in three independent biological replicates.

**Supplementary Figure S4. ChIRP-seq quality control and peak annotation for *lnc-ARRDC4-1*.** (A) Validation of lnc-ARRDC4-1 enrichment in three independent biological replicates of ChIRP-seq libraries by qPCR. The bars indicate the percentage of adjusted input recovered for *lnc-ARRDC4-1* and for the negative control transcripts *XIST* (libraries 1–2) or *MALAT1* (library 3), using total input (Input), the *lnc-ARRDC4-1*-specific probe pool (Even), and the nontargeting *LacZ* probe pool (*LacZ*). (B) PCA of ChIRP-seq signals across samples pulled down with *lnc-ARRDC4-1* and *LacZ* (control) probe sets, confirming separation between the experimental and control conditions. (C) Genomic feature distribution of high-confidence *lnc-ARRDC4-1* ChIRP-seq peaks (score ≥ 125). (D) Distribution of *lnc-ARRDC4-1* ChIRP-seq target genes (19171) by gene biotype. This experiment was performed in three independent biological replicates; the *q*-values < 0.05 were calculated using the Benjamini–Hochberg method.

**Supplementary Figure S5. The level of *DHX36* mRNA expression.** RT‒qPCR analysis of *DHX36* mRNA expression levels in lnc-ARRDC4-1 OE cells and lnc-ADCYAP1-2 OE cells was done relative to those in HEK WT cells. *GAPDH* levels were used for normalization. The data are presented as the mean ± SD (*n* = 4); *p*-values were calculated using two-sided Student’s *t* test (\**p*-value < 0.05; \*\**p*-value < 0.01; and \*\*\**p*-value < 0.001).

**Supplementary Figure S6. Polysome profiling of HEK WT, lnc-ADCYAP1-2 OE and lnc-ARRDC4-1 OE cells.** Polysome profiles measured by absorbance at 254 nm across 15–60% sucrose gradient fractions in HEK WT (black), lnc-ADCYAP1-2 OE (red) and lnc-ARRDC4-1 OE (blue) cells. Positions of the 40S and 60S ribosomal subunits and the 80S monosomes, light polysomes and heavy polysomes are indicated. This experiment was performed in three independent biological replicates.

**Supplementary Figure S7. Proteomic analysis of DHX36 overexpression in lnc-ARRDC4-1 OE and lnc-ADCYAP1-2 OE cells.** (A) Western blot validation of DHX36 overexpression. Representative immunoblots of DHX36 and TUBB in HEK WT cells and in lnc-ARRDC4-1 OE or lnc-ADCYAP1-2 OE cells transfected with EV or DHX36 OE (left panel). Quantification of DHX36 protein levels normalized to those of TUBB. The data are presented as the mean ± SD (*n* = 3); the *p*-values were calculated using two-sided Student’s *t* test (\**p*-value < 0.05 and \*\**p*-value < 0.01) (right panel). (B) PCA of mass spectrometry data from lnc-ARRDC4-1 OE cells transfected with empty vector (EV) and lnc-ARRDC4-1 OE cells transfected with the DHX36 overexpression vector (DHX36 OE). (C) PCA of mass spectrometry data from lnc-ADCYAP1-2 OE cells transfected with EV and lnc-ADCYAP1-2 OE cells transfected with DHX36 OE. (D) PCA under all conditions: HEK WT cells transfected with EV, lnc-ARRDC4-1 OE cells transfected with EV, lnc-ARRDC4-1 OE cells transfected with DHX36 OE, lnc-ADCYAP1-2 OE cells transfected with EV, and lnc-ADCYAP1-2 OE cells transfected with DHX36 OE. (E) Volcano plot of the differentially expressed proteins in lnc-ARRDC4-1 OE cells transfected with DHX36 OE compared with lnc-ARRDC4-1 OE cells transfected with EV. (G) Volcano plot of the differentially expressed proteins in lnc-ADCYAP1-2 OE cells transfected with DHX36 OE compared with lnc-ADCYAP1-2 OE cells transfected with EV. This experiment was performed in three independent biological replicates; the *q*-values < 0.05 were calculated using Welch’s *t* test in Perseus, and permutation-based FDR correction was applied for multiple testing.

**Supplementary Figure S8. Cellular localization of *lnc-ADCYAP1-2*.** Subcellular fractionation analysis of the distribution of *lnc-ADCYAP1-2* across chromatin, nucleoplasmic, and cytoplasmic fractions in lnc-ADCYAP1-2 OE cells, shown as chromatin-to-cytoplasm, chromatin-to-nucleoplasm, and nucleoplasm-to-cytoplasm ratios. *GAPDH* and *Actin* were used as cytoplasmic controls; *MALAT1, KCNQ10T1* and U1 snRNA were used as nuclear/chromatin controls. The data are presented as the mean ± SD (*n* = 4); the *p*-values were calculated using two-sided Student’s *t*-tests (*p*-value < 0.05; \*\**p*-value < 0.01; and \*\*\**p*-value < 0.001).

