## Supplementary figures for "Waking the sleepers: lincRNA overexpression compromises DHX36 activity and global protein synthesis"

### Supplementary Figure S1

#### Bulk tissue gene expression for ENSG00000259870 - *lnc-ARRDC4-1*

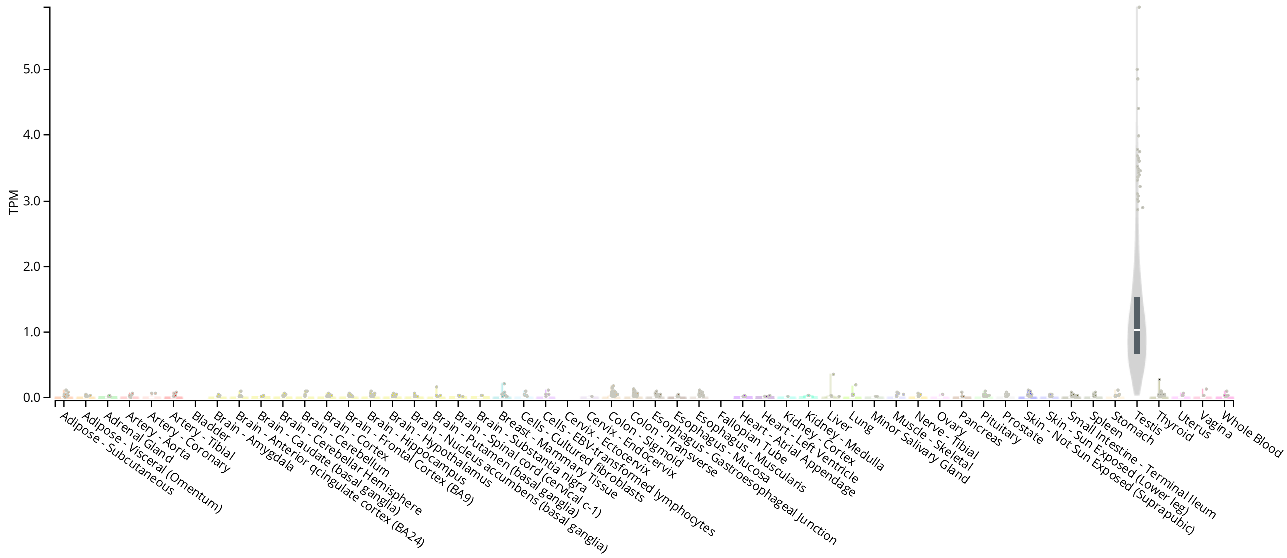


#### Bulk tissue gene expression for ENSG00000263551 - *lnc-ADCYAP1-2*

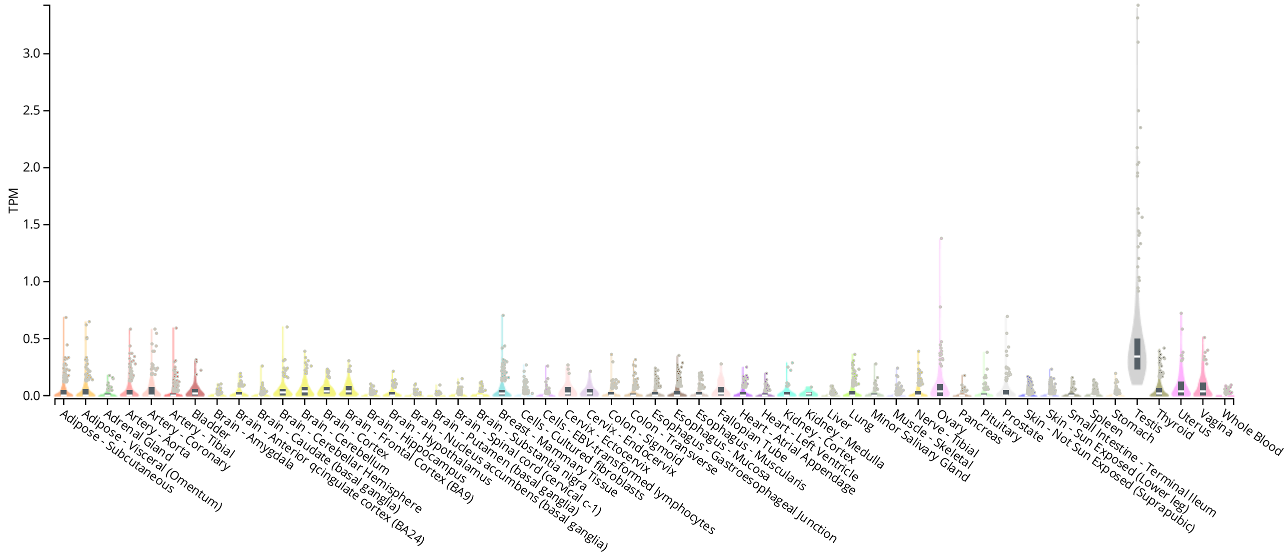

### Supplementary Figure S2

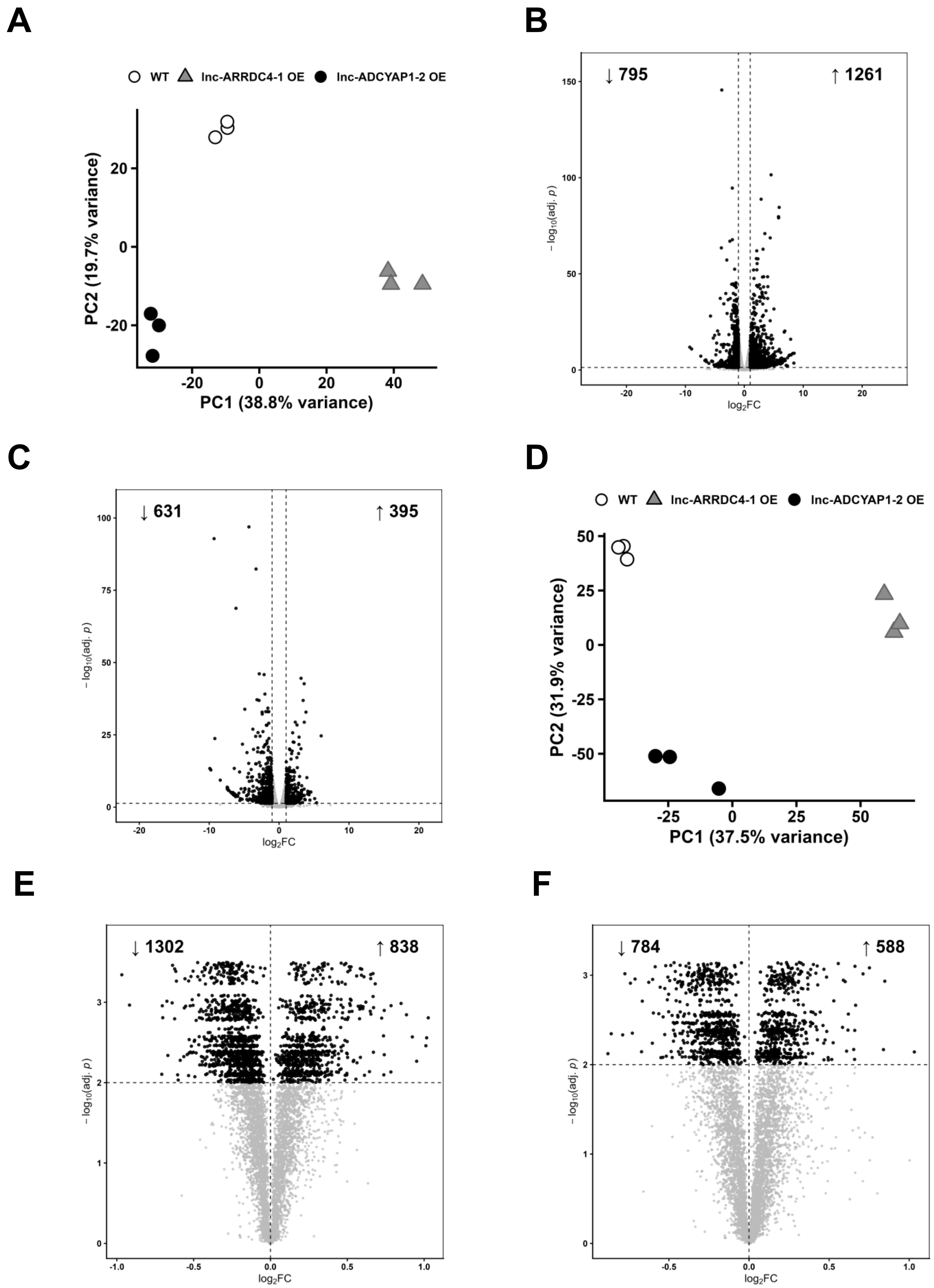

### Supplementary Figure S3

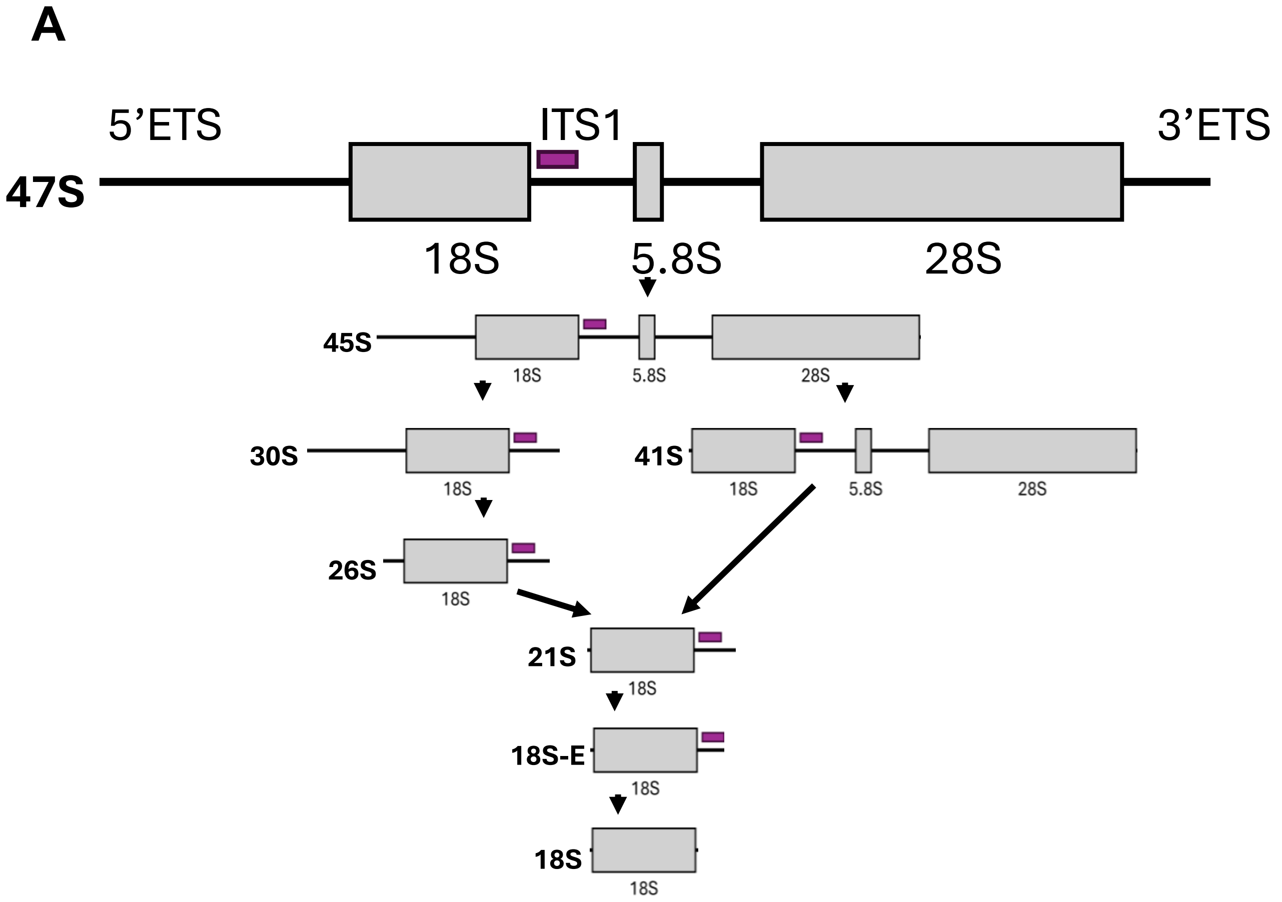


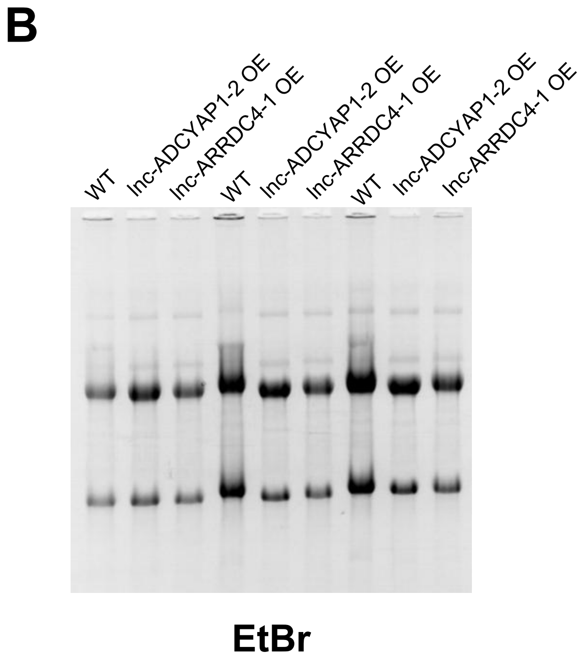


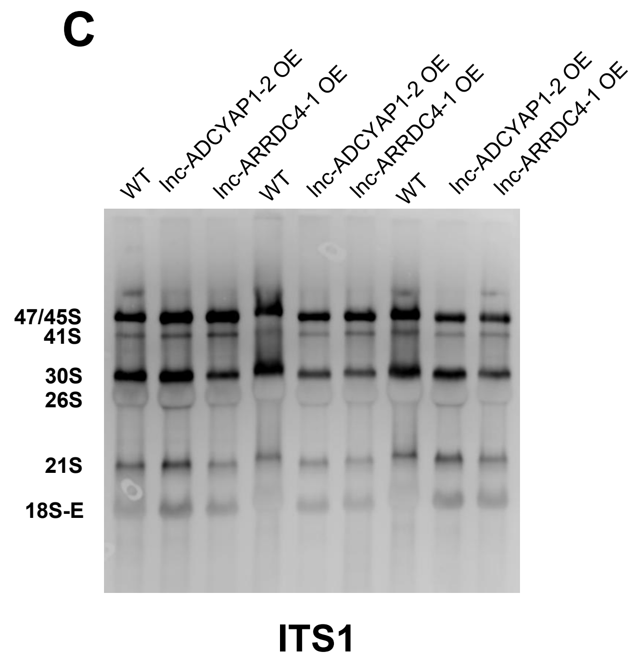

#### Supplementary Figure S4

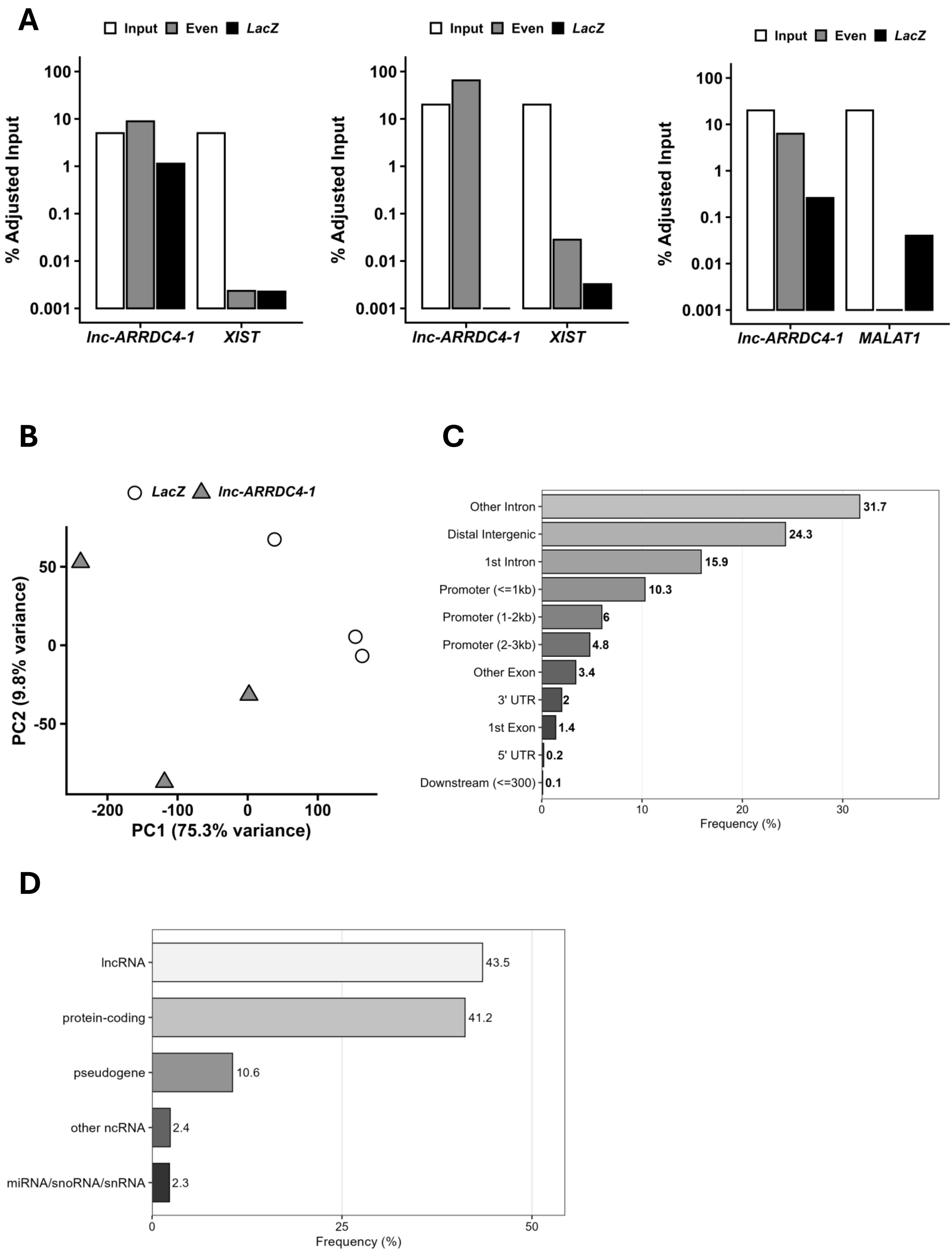

**Supplementary Figure S5**

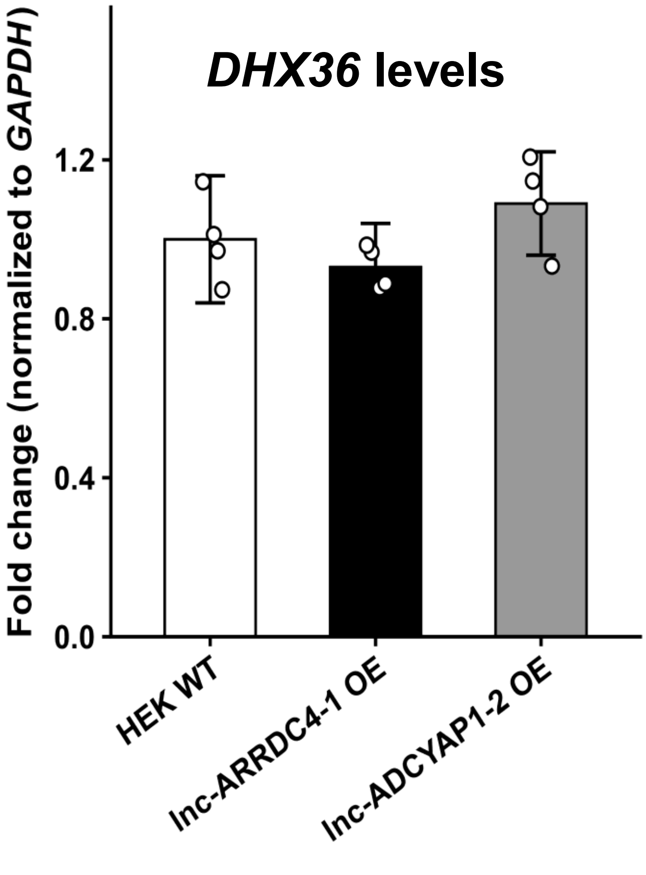

### Supplementary Figure S6

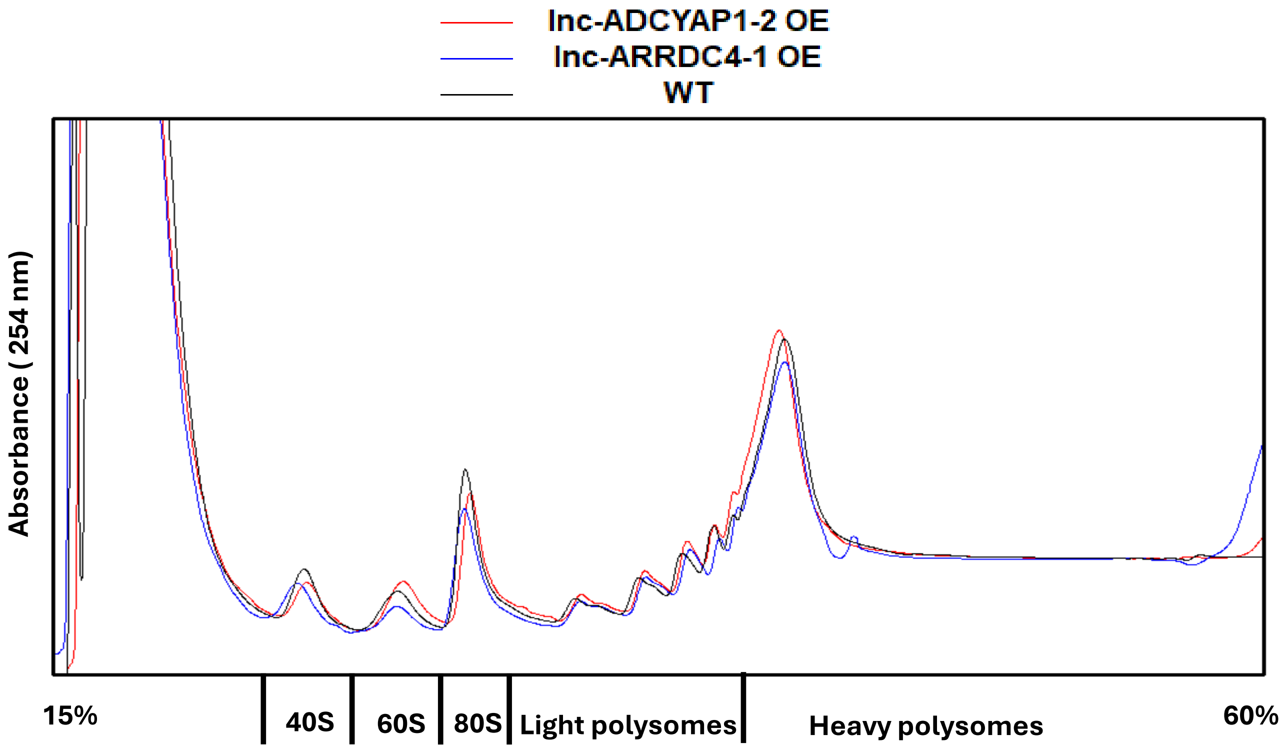

### Supplementary Figure S7

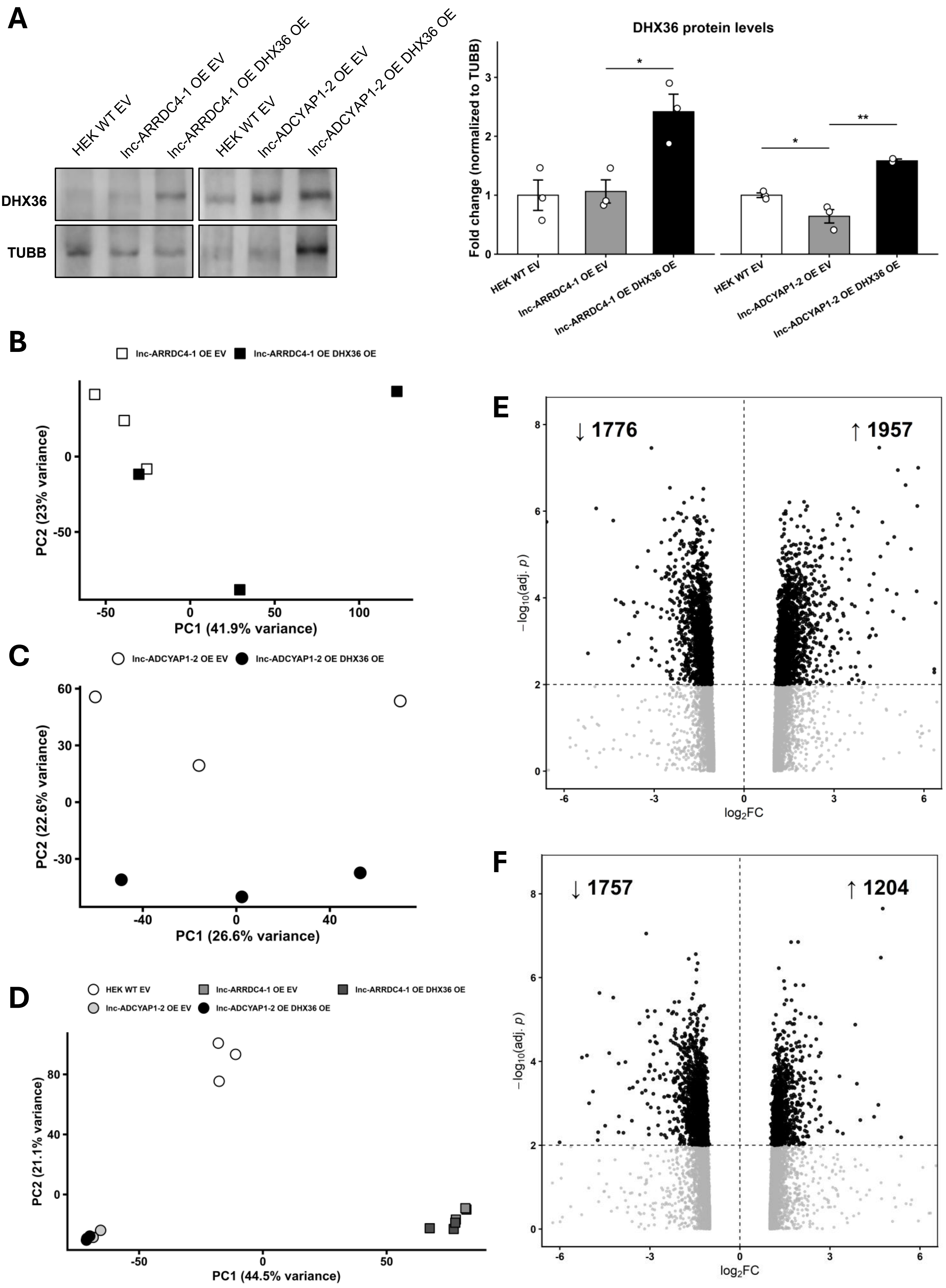

### Supplementary Figure S8

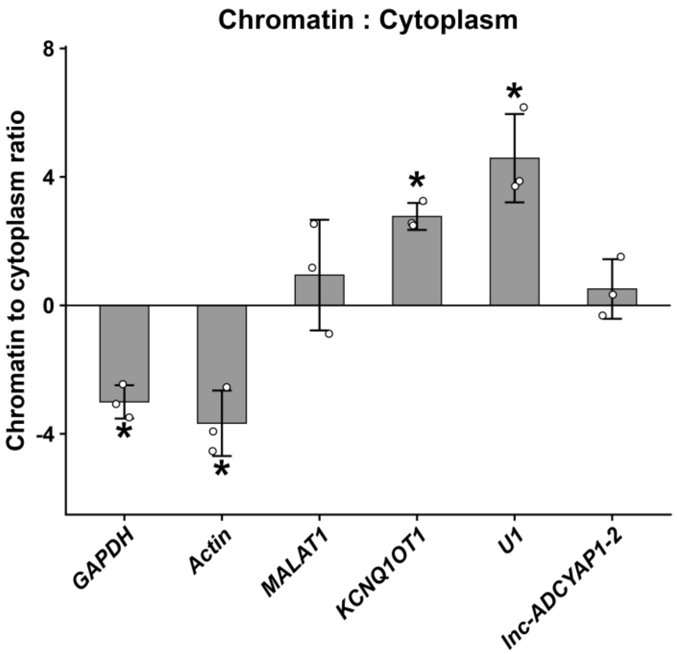


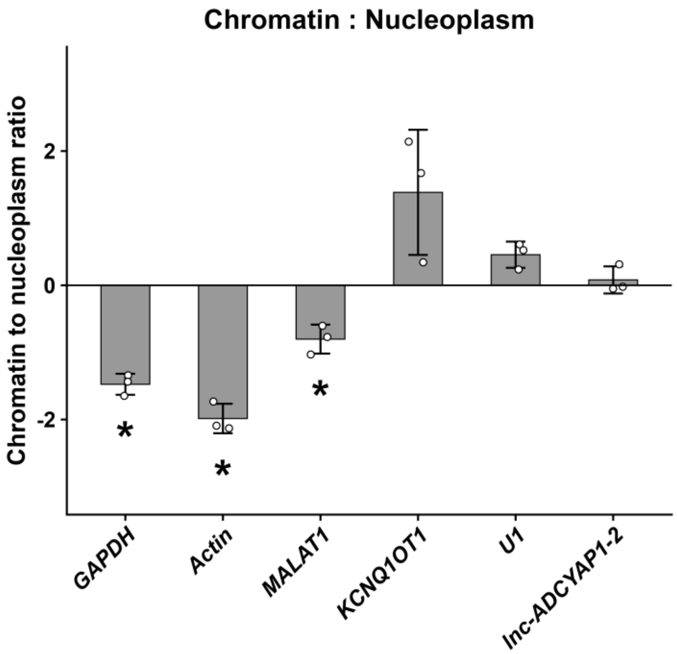


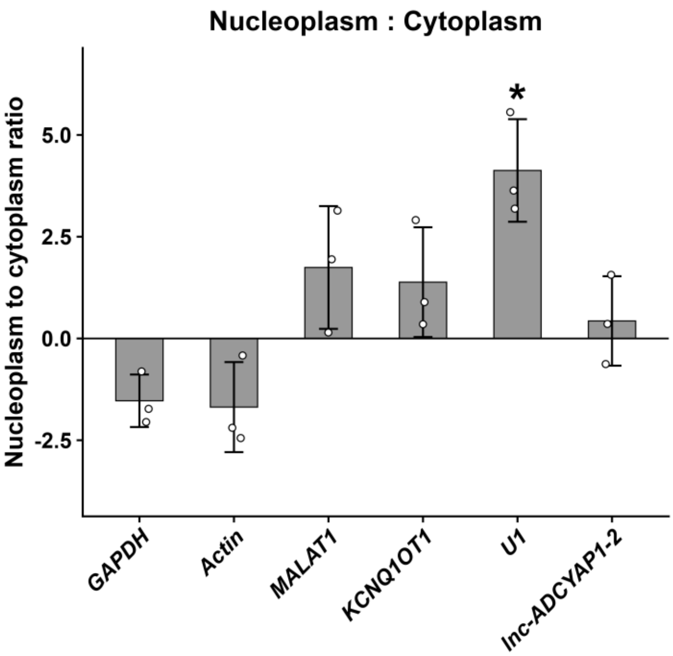
